# The Role of Fibrinogen-Mediated Platelet Aggregation in Subsequent Platelet-Driven Blood Clot Contraction

**DOI:** 10.64898/2026.06.23.733888

**Authors:** Alina I. Khabirova, Rafael R. Khismatullin, Shakhnoza M. Saliakhutdinova, Natalia G. Evtugina, Lorena Buitrago, Prashant K. Purohit, Rustem I. Litvinov, John W. Weisel

## Abstract

**Background:** Blood clot contraction/retraction depends on the force-generating actomyosin and on the platelet integrin αIIbβ3, which transmits intracellular forces to fibrin. Before clotting, fibrinogen binds to activated integrin αIIbβ3, mediating platelet aggregation. The relationship between platelet aggregation and subsequent platelet-driven clot contraction remains unclear.

**Methods:** We investigated the effects of platelet aggregation on clot contraction by selectively blocking the αIIbβ3-fibrinogen binding using the RGDW peptide. The ability of RGDW to disrupt αIIbβ3-fibrinogen binding was assessed by platelet aggregometry. The time-course of clot contraction was monitored optically in whole blood or platelet-rich plasma and modeled mathematically. Clot stiffness was assessed using Thromboelastography. The effect of the RGDW peptide on the structure of PRP-clots was examined using scanning electron microscopy.

**Results:** The RGDW peptide dose-dependently inhibited TRAP-induced platelet aggregation. Both in whole blood and in plasma, the peptide dose-dependently prolonged the lag-period and slowed the rate without affecting the final extent of contraction. Thromboelastography showed that RGDW dose-dependently increased maximum clot stiffness in blood. Scanning electron microscopy revealed that RGDW treatment resulted in formation of smaller fibrin agglomerates surrounding non-aggregated platelets. A theoretical model allowed us to decipher mechanisms underlying the kinetic effects of RGDW.

**Conclusion:** Blocking the binding of integrin αIIbβ3 to fibrinogen and preventing platelet aggregation delays and slows subsequent clot contraction without affecting the final degree of shrinkage. These findings indicate a modulatory role of fibrinogen-mediated platelet aggregation in clot contraction and highlight the unforeseen effects of selective inhibitors of platelet aggregation on the contraction of blood clots and thrombi.

## Introduction

Platelets are anuclear blood cells measuring 2-4 µm that are formed from megakaryocytes and circulate in the bloodstream for 7-10 days.^1^ Platelets play a key role in hemostasis and thrombosis due to a number of established reactions: i) adhesion to sub-endothelial structures at the site of vascular injury; ii) formation of platelet aggregates; iii) secretion of various bioactive substances, and iv) the procoagulant activity of the cell membrane, on which active enzyme complexes of blood coagulation factors are assembled. ^1^

In recent years, platelets have been shown to have another important biological property *in vivo*, namely their ability to cause contraction (compression or retraction) of hemostatic blood clots and obstructive thrombi.^2^ Contraction of blood clots and thrombi occurs as a result of traction forces generated by activated platelets attached to fibrin fibers, which form the 3D mechanical framework and transfer the contractile forces throughout the blood clot or thrombus. Platelet contractility is based on the ATP-dependent interaction of non-muscle myosin IIA with actin filaments, which leads to the generation of mechanical forces ranging from 1.5 to 79 nN per one platelet.^3^ Intracellular cytoskeletal contraction is transmitted via actin filaments across the membrane to the extracellular fibrin network, leading to its compaction, structural remodeling and a decrease in the volume of the entire clot due to extrusion of liquid blood serum.^2^

Blood clot contraction *in vivo* has important pathophysiological significance for both hemostasis and thrombosis. Compaction of the hemostatic clot brings the wound edges together and creates an impermeable seal, improving hemostasis and creating conditions for subsequent healing.^4,5^ Compression and compaction of the intravascular thrombus reduces occlusion and increases the size of the vessel lumen, thereby improving local blood flow.^6^ It also reduces the risk of thrombus rupture and embolization^7,8^ and affects its sensitivity to internal and external fibrinolysis.^9,10^

In addition to the platelet contractile machinery, the adhesive surface receptor integrin αIIbβ3 plays a critical role in blood clot contraction, serving as a mechanical bridge between the intracellular actin cytoskeleton and extracellular fibrin.^11^ On the one hand, the integrin αIIbβ3 is involved in mechanotransmission, i.e., the transmission of intracellular contractile forces to the extracellular fibrin matrix. On the other hand, the interaction of the integrin αIIbβ3 with fibrin causes the activation of intracellular signaling pathways depending on the elasticity of fibrin, a process known as mechanosensing that modulates the functional state of platelets, including contractility.^12^ Mechanotransmission and mechanosensing are oppositely directed processes of cellular mechanotransduction.^13–15^

Integrin αIIbβ3 is the most abundant adhesion receptor on the platelet surface, represented by 50,000–100,000 copies per platelet.^16^ It is a heterodimer consisting of α- and β-subunits that form an extracellular domain, a transmembrane domain, and a cytoplasmic domain linked to actin via adaptor proteins.^17,18^ In resting platelets, integrin molecules are essentially inactive and exist in a bent conformational state characterized by low affinity for ligands. Platelet activation, regardless of the nature of the stimulus, is accompanied by conformational extension of integrin, which acquires high affinity for its favorite ligands, fibrinogen and fibrin.^19,20^

The well-studied function of integrin αIIbβ3 is its participation in platelet aggregation mediated by the soluble plasma protein fibrinogen, which has a rod-like shape (45 nm by 3-6 nm).^2^ Activated integrin on the cell membrane binds to the lateral globular γ-nodules of fibrinogen, which has two binding sites for integrins and forms a bridge between adjacent platelets, leading to the formation of platelet aggregates.^21^ Interaction with integrin is mediated by the AGDV motif in the C-terminus of the γ-chain of human fibrinogen and can be inhibited by RGD-containing peptides.^19, 22^

However, activated αIIbβ3 also interacts with insoluble fibrin fibers, formed as a result of the enzymatic conversion of fibrinogen to monomeric fibrin followed by spontaneous polymerization. The interaction of integrin αIIbβ3 with fibrin differs from the interaction with fibrinogen in specificity and strength, since additional binding sites for integrin αIIbβ3, which were cryptic in the fibrinogen molecule, are exposed in fibrin.^23,24^ Binding of integrin αIIbβ3 to fibrin fibers ensures mechanotransmission and, ultimately, contraction of blood clots under the action of activated platelets.^15, 25^

Thus, integrin αIIbβ3 is involved in both platelet aggregation (mostly via fibrinogen) and clot contraction (via binding to fibrin). Both processes occur during blood coagulation, although aggregation of thrombin-activated platelets most likely begins earlier than clot formation and contraction. Obviously, these two processes, aggregation and contraction, which require the participation of active integrin αIIbβ3, are closely linked. However, it remains unclear how exactly the initial binding of activated platelets to fibrin(ogen) and their aggregation influence subsequent clot contraction. Is platelet aggregation necessary for subsequent clot contraction? Can platelet aggregation modulate the kinetics and extent of subsequent clot contraction? Do platelet aggregates differ from single activated platelets in the total area of interaction with fibrin fibers, in the contractile force generated, and in the ability to change the structure and properties of the fibrin network and the entire blood clot? These questions are not only of theoretical importance for understanding the functional diversity and plasticity of platelets but also have clinical implications in connection with the development of antiplatelet drugs capable of selectively modifying individual platelet functions.^26–29^

Clearly, to answer these questions experimentally, it is necessary to separate platelet aggregation and clot contraction, i.e., to study contraction under conditions where platelet aggregation is specifically suppressed by disrupting the interaction of integrin αIIbβ3 with fibrinogen while maintaining its binding to fibrin. We selected the RGDW (Arg-Gly-Asp-Trp) peptide as a tool for studying this problem. The amino acid sequence RGD (Arg-Gly-Asp) is known to be recognized by many integrins, including αIIbβ3, and is present in protein ligands such as fibrinogen, fibronectin, and vitronectin.^30^ As shown by Buitrago et al. (2021),^31^ the RGDW peptide, which contains the additional tryptophan, is capable of selectively blocking fibrinogen binding to integrin without affecting its interaction with fibrin. Specifically, RGDW was shown to inhibit the initial phase of fibrinogen-mediated platelet aggregation but did not affect the second wave of light transmission caused by platelet interaction with polymerizing fibrin. These results provide evidence that platelet–fibrin interactions involved in clot contraction are mediated by a mechanism distinct from that governing the interaction between αIIbβ3 and fibrinogen.

Based on the above, the aim of this work was to study the effect of platelet aggregation on the contraction of blood clots by selectively blocking the interaction of integrin αIIbβ3 with fibrinogen under the action of the RGDW peptide. The results show that RGDW delays clot contraction and shifts clot morphology toward the formation of small platelet-fibrin agglomerates composed of individual platelets. The practical significance of these findings lies in the potential for developing agents that selectively modulate platelet functions to alter the kinetics of clotting and clot or thrombus structure.

## Materials and Methods

### Experimental materials and methods

Details on the collection and fractionation of human blood samples, light transmission platelet aggregometry, clot contraction assay, Thromboelastography, scanning electron microscopy (SEM) of platelets and quantitative SEM image analysis, as well as statistical analyses, are provided in Supplemental Material, Section 1.

### Theoretical model

To get mechanistic insights, a mathematical model was constructed to track as a function of time the populations of platelets that were quiescent and activated, the latter bound to fibrinogen, fibrin or the RGDW peptide. The model consists of series of ordinary differential equations for these populations that were integrated numerically in MATLAB (see Supplemental Material, Section 2, for details). The model assumes that (a) fibrin and fibrinogen once bound to platelets are not released, (b) RGDW can bind and unbind from platelets, (c) platelets bound to RGDW can bind to fibrin, but not fibrinogen, (d) platelet activation and aggregation involves positive feedback since aggregated platelets release compounds (ADP, TxA_2_) that activate other platelets, and (e) activated platelets bound to fibrin exert forces on the poroelastic fibrin network that cause it to shrink. The rate constants associated with the binding of platelets to fibrin, fibrinogen and RGDW are left as fitting parameters that are adjusted to reproduce the trends, extent of clot contraction and time to full contraction seen in experiments. Importantly, one set of rate constants captures all trends in clot contraction as a function of time irrespective of the RGDW concentration allowing us to make predictions.

## Results

### Selection of working concentrations of the RGDW peptide based on the ability to inhibit platelet aggregation

To quantify the ability of the RGDW peptide to inhibit the interaction of platelet integrin αIIbβ3 with fibrinogen, TRAP-induced platelet aggregation was used. RGDW peptide at increasing final concentrations (50, 100, 200, and 300 μM) was incubated with platelet-rich plasma for 5 minutes at 37°C before adding the aggregation trigger.

As shown in Fig. 1, the final degree of aggregation induced by 10 μM TRAP without the addition of RGDW peptide (positive control) averaged 58±4%. The addition of RGDW caused a significant dose-dependent inhibition of aggregation (p<0.0001, one-way ANOVA). Thus, at the minimum concentration of RGDW peptide used (50 μM), a slight decrease in the degree of final aggregation to 51±6% was observed, which was not statistically significant compared to the control (p=0.4). Starting from a concentration of 100 μM and more, the RGDW peptide had a significant dose-dependent anti-aggregation effect. At 100 μM of the peptide, the average final degree of aggregation decreased to 39±4% (p=0.02), at 200 μM the final degree of aggregation was 21±4% (p=0.02), and at 300 μM the degree of aggregation decreased to 10±2% (p=0.02). Thus, under these experimental conditions, the RGDW peptide effectively and dose-dependently suppresses TRAP-induced platelet aggregation in the concentration range of 100–300 μM, although the first signs of inhibition appear already at a concentration of 50 μM.

**Figure 1.**
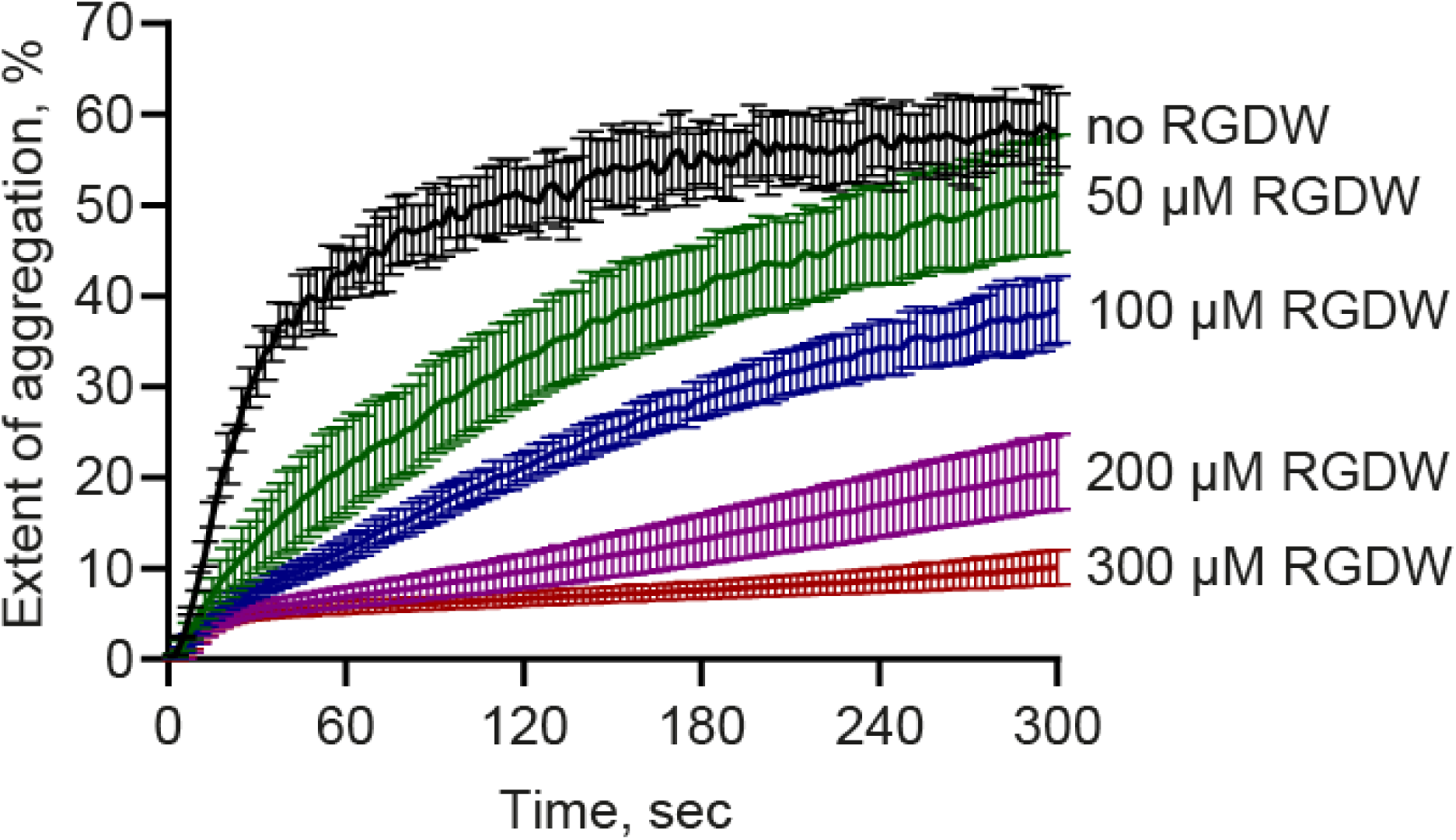
Platelet aggregation in PRP induced by 10 μM TRAP in the absence (*black curve*) and presence of the RGDW peptide at various concentrations of 50 μM, 100 μM, 200 μM, 300 μM. The results are presented as the mean ± SEM (n=3). For numerical data see Table S1.

### Effects of the RGDW peptide on clot contraction in whole blood

To assess the effects of the RGDW peptide on the ability of platelets to induce clot contraction, the kinetics of thrombin-induced clot contraction was studied. The results are presented in Fig. 2.

**Figure 2.**
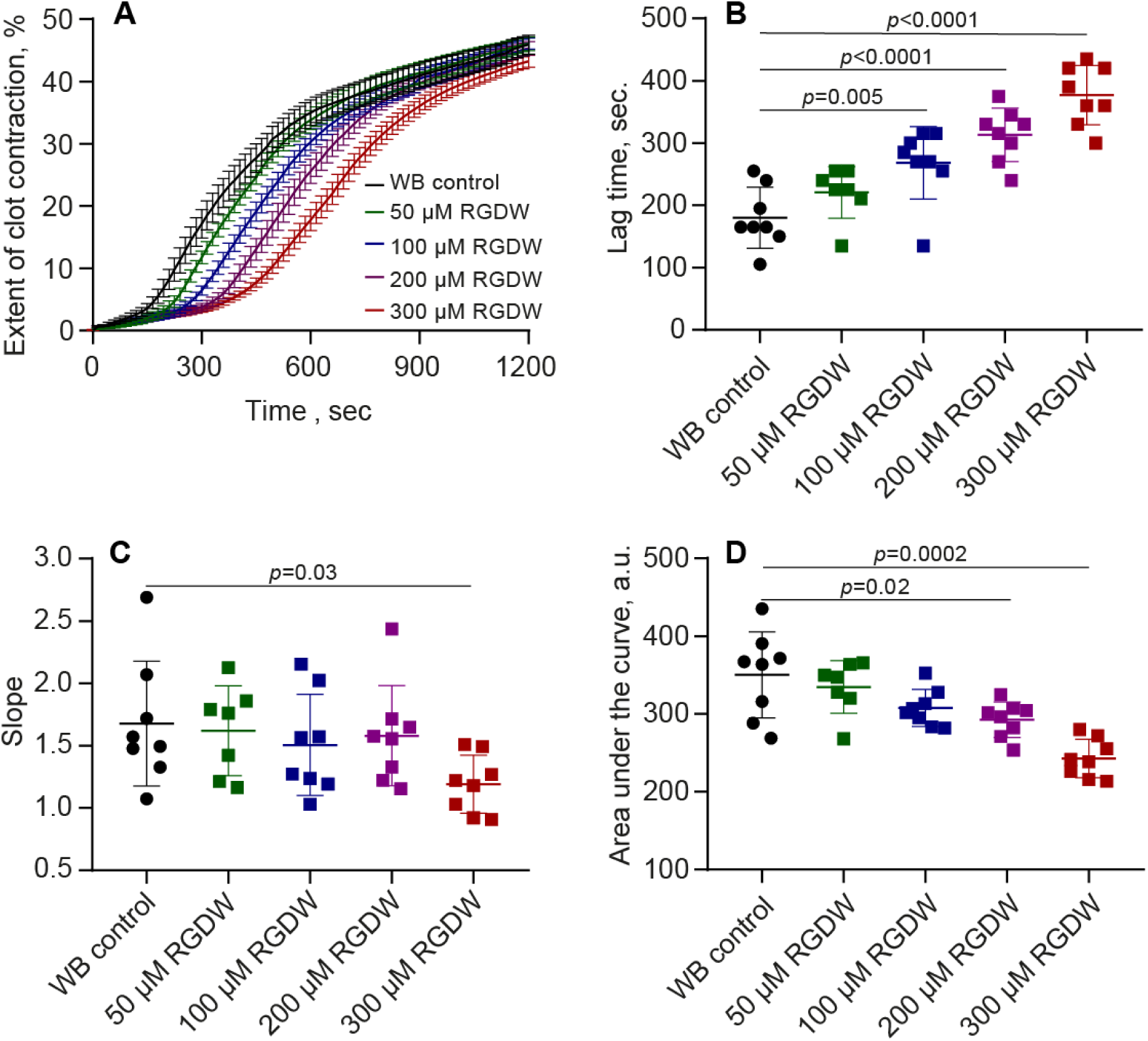
Contraction kinetic curves (**A**) and contraction parameters (**B-D**) of **clots formed in whole blood** (WB) in the absence and presence of the RGDW peptide at various concentrations of 50 μM, 100 μM, 200 μM, 300 μM. The results are presented as the mean ± SEM (n=7-8). The *p*-values reflect significant differences with control determined with the unpaired Student’s *t*-test. For numerical data see Table S3.

The addition of RGDW peptide in the concentration range of 50–300 μM did not have any noticeable effect on the final size of the contracted clot. The final extent of clot contraction in the control sample without RGDW averaged 46±2% and remained virtually unchanged (46±1%) at peptide concentrations of 50, 100, and 200 μM. Only at the highest concentration used, 300 μM, there was a slight decrease in the final extent of contraction to 43±1%, which, however, did not reach the level of statistical significance (p=0.5). The overall analysis of concentration dependence (one-way ANOVA) revealed no significant differences between the groups of data (p=0.3).

In contrast to the final extent of contraction, the RGDW peptide exerted a dose-dependent inhibitory effect on the kinetic parameters of clot contraction - the onset time or lag period, rate (slope of the kinetic curve) and total intensity of the process (area under the kinetic curve), although the evidence of this effect was different. Thus, the average lag time in the control samples (without RGDW) was 180±17 sec.; at concentrations of 50, 100, 200 and 300 μM this value increased to 221±16 sec. (p=0.02), 268±21 sec. (p=0.01), 313±15 sec. (p=0.001) and 377±17 sec. (p=0.001), respectively. Analysis of variance confirmed a statistically significant dose-dependent effect of RGDW on the lag time (p<0.0001).

The average slope of the kinetic curves in the control samples (without RGDW) was 1.68±0.18. RGDW peptide at concentrations of 50, 100, and 200 μM did not have a statistically significant effect on this parameter, the values of which were in the range of 1.51–1.62 (p>0.05 for all data compared to the control). Only at the highest concentration studied (300 μM) there was a significant decrease in the average slope to 1.19±0.08 (p=0.03). The overall trend, assessed using analysis of variance, did not reach the level of statistical significance (p=0.1).

The RGDW peptide had a clear dose-dependent effect on the average area under the kinetic curves, which in the control (without RGDW) was 350±20 a.u. Addition of the peptide at concentrations of 50 and 100 μM led to a decrease in this parameter at the border of statistical significance to 335±13 a.u. (p=0.06) and 308±8 a.u. (p=0.055), and at concentrations of 200 and 300 μM, the decrease in the area under the kinetic curve was significant (293±8 a.u., p=0.02 and 243±9 a.u., p=0.001, respectively). The overall dynamics of the decrease in the area under the kinetic curve was also highly significant (p<0.0001, one-way ANOVA).

Thus, in whole blood, the RGDW peptide at the concentrations studied causes a dose-dependent and statistically significant slowdown in the onset of contraction (lag time) and a decrease in the overall intensity of clot contraction (area under the kinetic curve), exerting a moderate effect on the rate of the process (angle of inclination of the kinetic curve) and without changing the final extent of clot contraction.

For comparison, contraction of whole blood clots was studied in the presence of abciximab (brand name ReoPro™), a monoclonal antibody-based medication used to prevent blood clots during balloon angioplasty or stent placement. As a potent inhibitor of αIIbβ3, abciximab strongly suppressed platelet aggregation in a dose-dependent manner (Fig. S1). Abciximab also inhibited blood clot contraction, but, unlike RGDW, it reduced both the rate and final extent of clot contraction (Fig. S2), suggesting a broader anti-αIIbβ3 specificity.

### Effect of the RGDW peptide on clot contraction in platelet-rich plasma

To assess the effect of the RGDW peptide on platelet contractility, which causes compression and remodeling of fibrin clots in the absence of red blood cells and white blood cells, the contraction of thrombin-induced clots in PRP was studied. The results are presented in Fig. 3.

**Figure 3.**
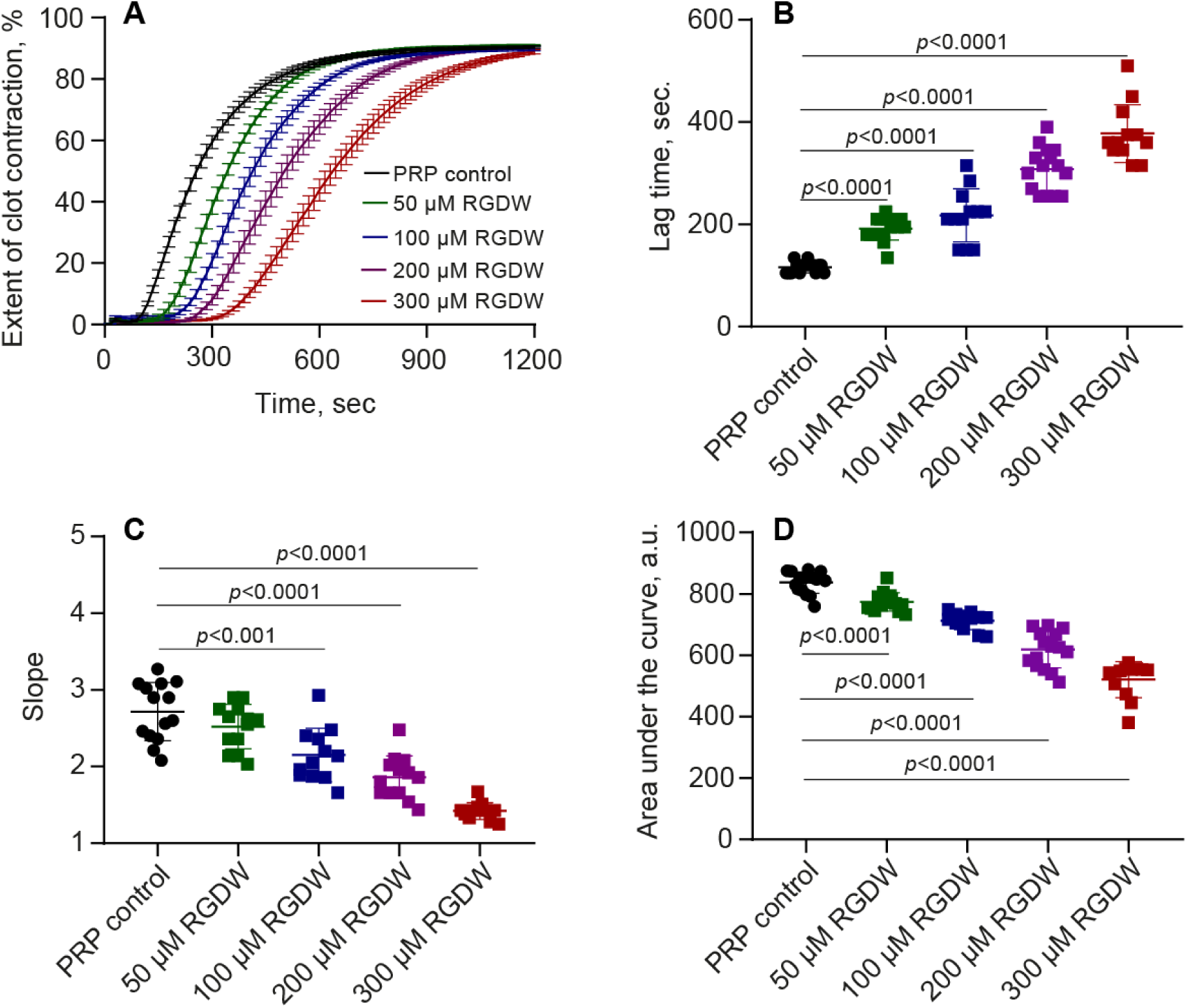
Contraction kinetic curves (**A**) and contraction parameters (**B-D**) of **clots formed in PRP** in the absence and presence of the RGDW peptide at various concentrations of 50 μM, 100 μM, 200 μM, 300 μM. The results are presented as the mean ± SEM (n=6-9). The *p*-values reflect significant differences with control determined with the unpaired Student’s *t*-test. For numerical data see Table S4.

As in whole blood, the addition of RGDW peptide in the concentration range of 50–300 μM did not affect significantly the mean final extent of clot contraction, which remained virtually constant within a narrow range of 89–91% in the absence and presence of RGDW. Similar to the whole blood data, in PRP the RGDW peptide caused a significant dose-dependent prolongation of the lag period, the time required for the onset of clot contraction. In the control (without RGDW), the average lag time was 122±9 sec, whereas at peptide concentrations of 50, 100, 200, and 300 μM, the lag time increased to 190±27 sec (p=0.02), 213±74 sec (p=0.01), 315±51 sec (p=0.001), and 398±78 sec (p=0.001), respectively. Analysis of variance confirmed the high statistical significance of the dose-dependent effect of RGDW on the lag time (p<0.0001).

The RGDW peptide exerted a pronounced dose-dependent inhibitory effect on the slope of the kinetic curves. In the control samples, the average slope tangent was 2.72±0.10. At peptide concentrations of 50, 100, 200, and 300 μM, a distinct dose-dependent decrease in this parameter was observed to 2.52±0.01 (p=0.15), 2.15±0.10 (p=0.001), 1.86±0.08 (p<0.0001), and 1.42±0.03 (p<0.0001), respectively. The overall dynamics of contraction rate decrease under the influence of increasing RGDW peptide concentrations was highly significant (p<0.0001, one-way ANOVA).

In accordance with the changes in the lag time and the contraction rate, the RGDW peptide had a pronounced dose-dependent effect on the area under the kinetic curve, which in the control (without RGDW) was 822±38 a.u. With increasing peptide concentrations of RGDW (50-300 μM), the area under the curve significantly decreased from 780±36 a.u. (p=0.02) to 498±77 a.u. (p=0.001). The overall pattern was statistically significant (p<0.0001, one-way ANOVA).

Thus, in PRP clots, the RGDW peptide dose-dependently inhibits the onset of contraction (increases the lag time), slows the process (the slope of the kinetic curves), and reduces the overall intensity of clot contraction (decreases the area under the kinetic curve), without affecting the final degree of clot shrinkage. The significantly more pronounced effect of the RGDW peptide on the contraction kinetics in plasma compared to whole blood can be explained by the fact that the absolute contraction rate of fibrin clots containing no red blood cells (∼3; Fig. 3C) is approximately 2 times higher than in whole blood (∼1.8; Fig. 2C), making the relative changes in this parameter more noticeable.

### Effect of RGDW peptide on clot elasticity in whole blood

To assess the effect of the RGDW peptide on the clot mechanical properties, Thromboelastography was used, in which blood clotting was induced by kaolin and calcium ions. The results are presented in Fig. 4.

**Figure 4.**
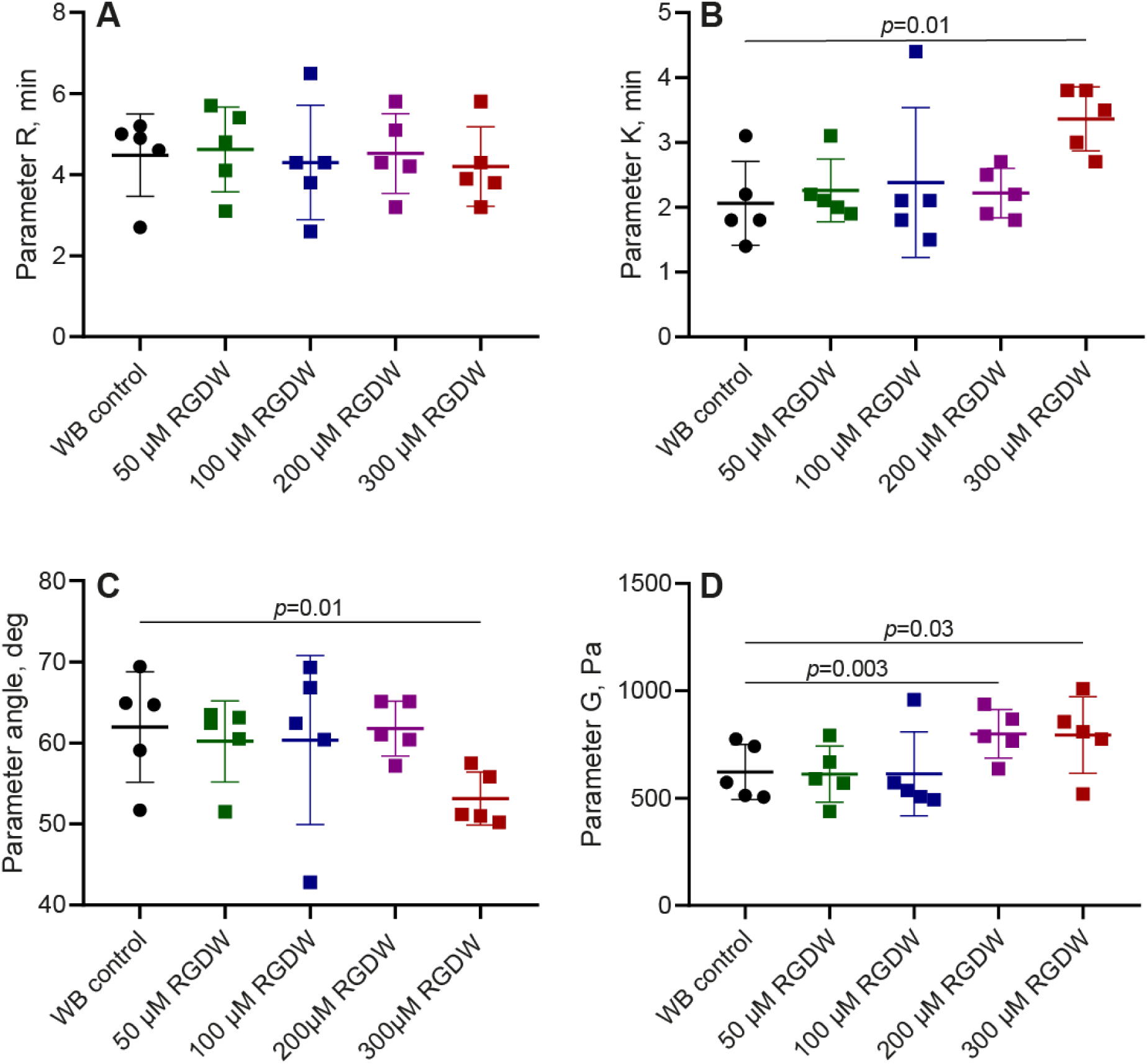
Parameters of TEG obtained in **whole blood** in the absence and presence of the RGDW peptide at various concentrations of 50 μM, 100 μM, 200 μM, 300 μM. The results are presented as the mean ± SEM (n=5). The *p*-values reflect significant differences with control determined with the Student’s paired *t*-test. For numerical data see Table S6.

RGDW at concentrations ranging from 50 to 200 μM did not significantly affect the time from initiation to onset of clotting (parameter *R*). The mean *R* values in the control (without RGDW) and experimental groups changed from 4.2±0.4 to 4.6±0.5 min without significant differences (p=0.5). Similarly, the clot formation rate, measured by the *K* parameter, averaged 2.1±0.3 min in the control and remained unchanged at the RGDW peptide concentrations ranging from 50 to 200 μM. Only at the maximum peptide concentration (300 μM) the *K* value increased significantly to 3.4±0.2 min (p=0.01 compared to the control), indicating a moderate slowdown in the clot formation rate. The overall trend toward an increase in the *K* parameter with increasing RGDW peptide concentration did not reach statistical significance (p=0.06, one-way ANOVA). The average *α* angle on the TEG, which, like the *K* parameter, characterizes growth of clot stiffness, also remained stable at peptide concentrations up to 200 μM, remaining within the control values (62°±3°). At a 300 μM RGDW peptide concentration, the *α* angle decreased to an average of 53°±1° (p=0.03), indicating a slowdown in blood clot formation. The overall dynamics of changes in the *α* angle as a function of the RGDW peptide concentration was insignificant (p=0.1, one-way ANOVA).

The RGDW peptide had the most pronounced effect on the mechanical properties of the final blood clot, assessed by the elastic modulus (*G*). In control samples (without RGDW), the average *G* value was 622±57 Pa. The RGDW peptide at concentrations of 50 and 100 μM did not affect this parameter. However, starting with a concentration of 200 μM, a significant increase in *G* was observed, reaching 800±51 Pa (p=0.01) at 200 μM and 794±80 Pa (p=0.03) at 300 μM. Analysis of variance confirmed the significance of the observed dose-dependent effect of RGDW peptide on blood clot elasticity (p=0.03, one-way ANOVA).

Thus, the RGDW peptide causes a dose-dependent increase in the maximum stiffness of the final blood clot, somewhat slows down the process of clot formation at relatively high concentrations, and has no visible effect on the time from initiation to the onset of blood clotting.

Compared to the RGDW peptide, abciximab had the opposite effect on the clot elasticity, reducing the clot stiffness in a dose-dependent manner (Fig. S3), again reflecting a broader specificity of the anti-αIIbβ3 activity that blocks platelet interactions both with soluble fibrinogen and insoluble polymerized fibrin, resulting in impaired clot contraction (Fig. S2).

### Effect of the RGDW peptide on clot elasticity in platelet-rich plasma

To assess the effect of the RGDW peptide on the dynamics of fibrin clot formation and clot mechanical properties in the absence of erythrocytes and leukocytes, Thromboelastography of kaolin-treated, re-calcified PRP was performed (Fig. 5).

**Figure 5.**
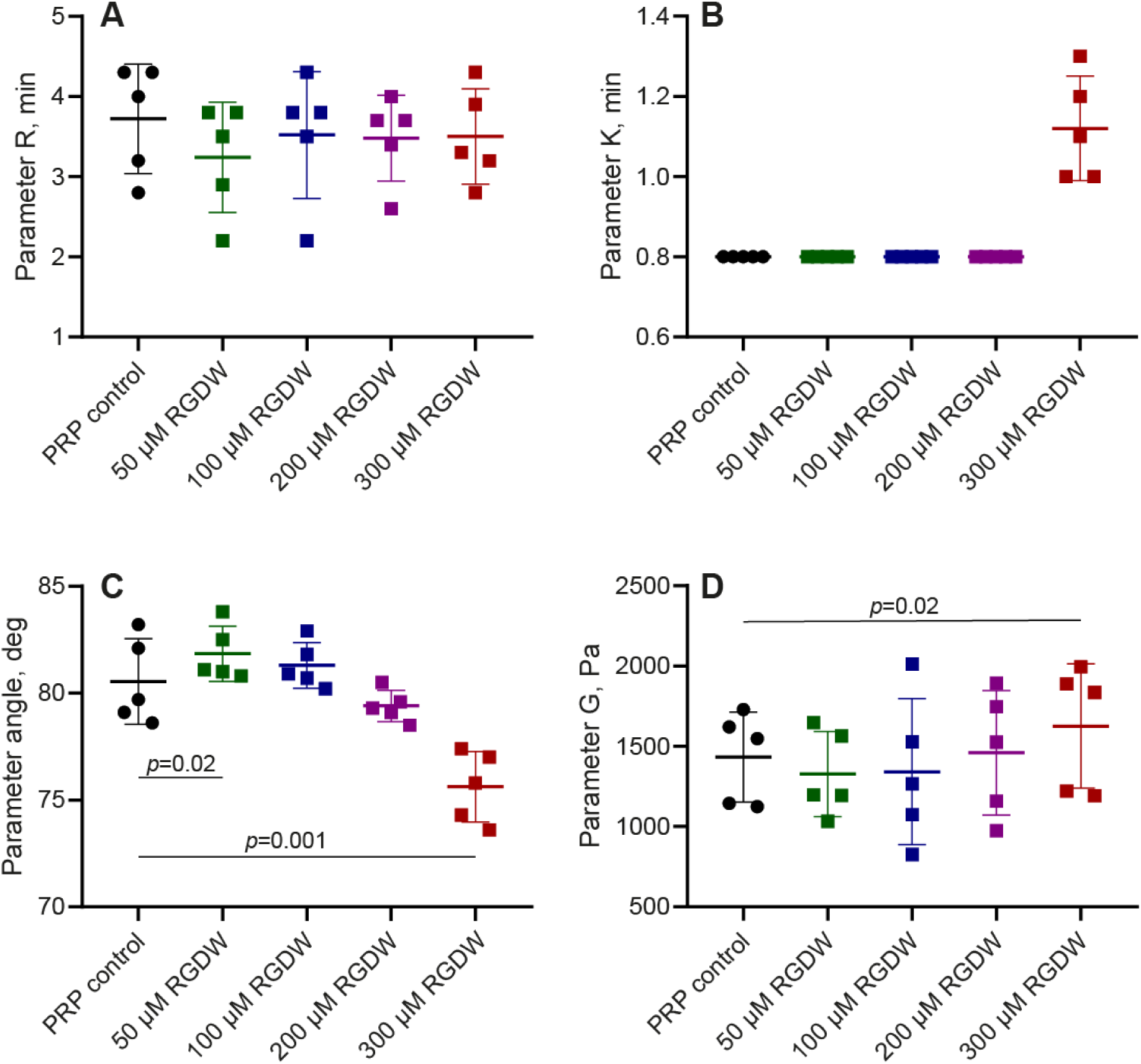
Parameters of TEG obtained in **PRP** in the absence and presence of the RGDW peptide at various concentrations of 50 μM, 100 μM, 200 μM, 300 μM. The results are presented as the mean ± SEM (n=5). The *p*-values reflect significant differences with control determined with the Student’s paired *t*-test. For numerical data see Table S7.

As in whole blood, the RGDW peptide did not significantly affect the time from initiation to onset of clotting (parameter *R*). The mean *R* values in the control (without RGDW) and in the experimental samples with different RGDW concentrations ranged from 3.2±0.3 to 3.7±0.3 min, with no significant differences (p=0.3, one-way ANOVA). The *K* parameter, which characterizes the rate of clot formation, also remained unchanged (0.80 ± 0.01 min) at peptide concentrations up to and including 200 μM. At the maximum peptide concentration (300 μM), a tendency toward an increase in *K* to 1.1±0.1 min was observed (p=0.06), indicating a moderate slowdown in the rate of clot formation and explaining the general pattern of a significant increase in *K* with increasing RGDW peptide concentration (p=0.05, one-way ANOVA).

The most pronounced dose-dependent inhibitory effect of the RGDW peptide on the rate of clot formation was observed based on changes in the *α* angle, which reflects the increase in the strength of the forming clot. At a peptide concentration of 50 μM, there was an insignificant rise in the mean value of the *α* angle to 82°±1° (p=0.2) compared to the control (81°±1°); however, with a further increase in the peptide concentration, the *α* angle progressively decreased to 79°±0.3° (p=0.02) at 200 μM and to 76°±1° (p=0.001) at 300 μM, indicating a dose-dependent slowing down of blood clot formation, confirmed by one-way analysis of variance (p<0.0001).

The effect of the RGDW peptide on the mechanical properties of the final clot in blood plasma had opposite directions depending on a concentration. Compared to the control (1430±125 Pa), a moderate decrease in the mean *G* values was observed at peptide concentrations of 50 μM (1330±119 Pa, p=0.5) and especially 100 μM (1340±203 Pa, p=0.4); a return to the control G values at a peptide concentration of 200 μM; and an increase at 300 μM (up to 1630±173 Pa, p=0.01). Not surprisingly, there was no overall dependence of clot elasticity on the RGDW peptide concentration (p=0.3, one-way ANOVA).

Thus, in PRP, the RGDW peptide exerts a pronounced dose-dependent inhibitory effect on the rate of fibrin polymerization (a decrease in the *α* angle), particularly noticeable at high peptide concentrations (200–300 μM). However, the peptide’s effect on the final clot stiffness (parameter *G*) varies across the concentrations applied. As in whole blood, the peptide has no significant effect on the time from initiation to onset of clotting in PRP.

The dose-dependent effect of abciximab on the decrease in clot elasticity in PRP (Fig. S4) was like that observed in whole blood (Fig. S3), reflecting weakening of the platelet-fibrin interactions, reduced mechanotransmission, and impaired clot contraction.

### Effect of the RGDW peptide on clot morphology in platelet-rich plasma

To evaluate the effect of the RGDW peptide on the structure of contracted clots formed by thrombin in PRP, we carried out scanning electron microscopy (SEM) followed by morphometric image analysis. Quantitative analysis included measuring the area of fibrin agglomerates formed around activated platelets.

In control (without RGDW) clots, fibrin formed a dense network of fibers, containing areas of irregularly shaped fibrin accumulation and compaction (Fig. 6A). These fibrin patches were interconnected by extended and stretched fibrin fibers or fiber bundles, where several fibers aggregated laterally to form thicker fibrils. The observed structure is characteristic of PRP clots with accumulations of compacted fibrin around activated platelets and their aggregates.^32^

**Figure 6.**
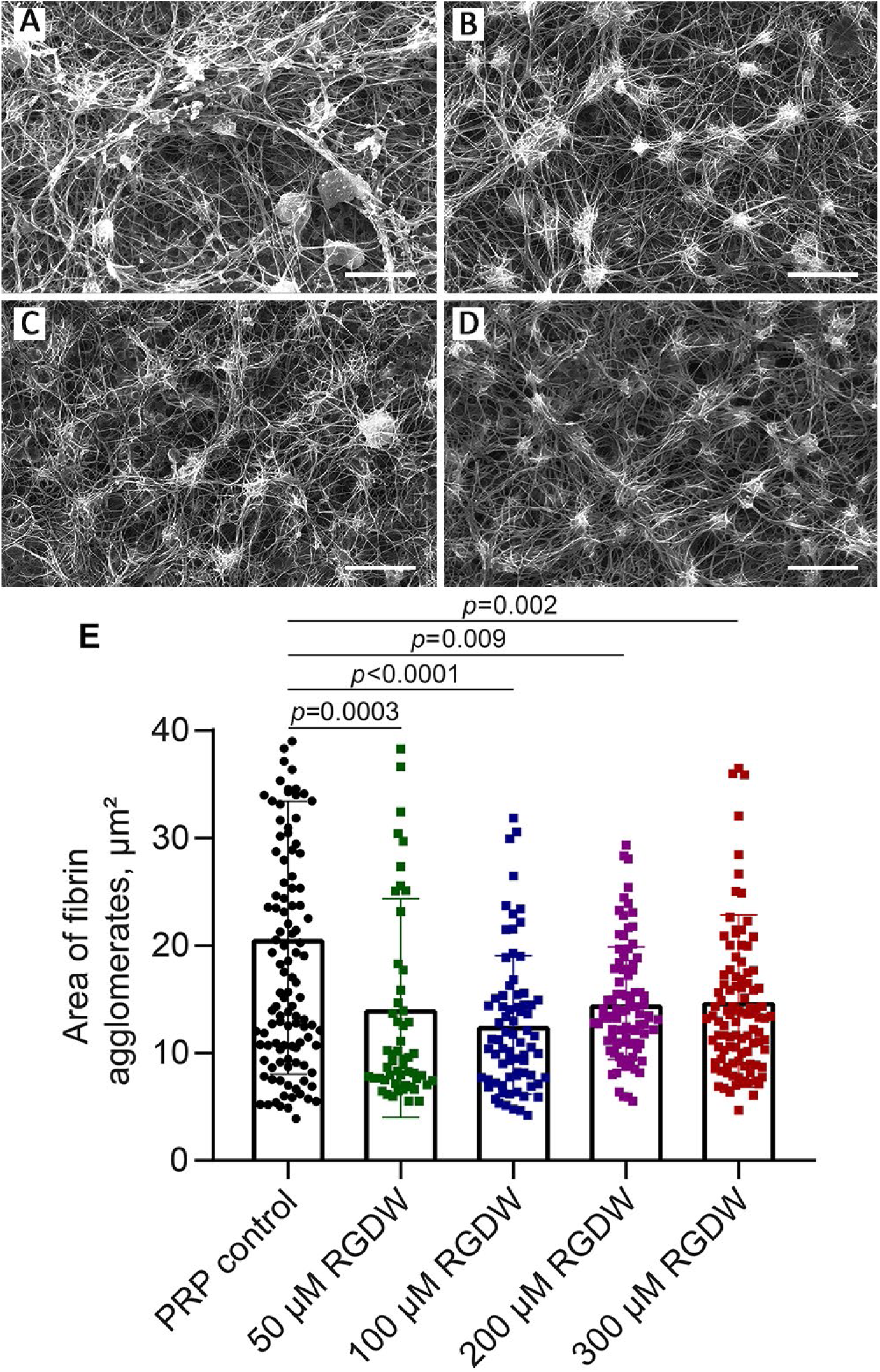
Representative scanning electron micrographs of contracted clots formed in PRP in the absence (**A**) and presence of the RGDW peptide at 100 μM (**B**), 200 μM (**C**), and 300 μM (**D**). Magnification bars = 10 μm. **(E)** The size (area) of fibrin agglomerates in contracted PRP clots formed in the absence and presence of the RGDW peptide at various concentrations of 50 μM, 100 μM, 200 μM, and 300 μM. Dots represent the sizes of individual fibrin agglomerates quantified in 5 random SEM images of clots obtained for each experimental condition and exemplified in A-D. The results are presented as the mean ± SEM. The *p*-values reflect the significant differences with PRP control determined using paired Student’s *t*-test. For numerical data see Table S10.

In clots formed and contracted in the presence of the RGDW peptide, the fibrin network had a more uniform, homogeneous structure with scattered fibrin agglomerated to a round-like or stellate shape (Fig. 6B-D). Fiber bundles were significantly less common, and the fibrin network itself consisted mainly of individual, wavy, unoriented fibers that were less clearly stretched between adjacent fibrin masses around platelets. Fibrin agglomerates were more frequent than in the control, but their size was visually smaller. This visible impression was confirmed by morphometric analysis (Fig. 6E). In control samples (without RGDW), the average area of fibrin agglomerates measured using ImageJ was 20.7±1.2 μm². The addition of the peptide resulted in a dose-dependent decrease in the size of fibrin agglomerates. Already at the minimum studied concentration (50 μM), the average area of fibrin agglomerates significantly decreased to 14.2±1.4 μm² (p=0.0003). The maximum effect was observed at a peptide concentration of 100 μM, when the area of agglomerates decreased to 12.6±0.8 μm² (p<0.0001). With a further increase in the peptide concentration (200 and 300 μM), the area of agglomerates still remained significantly smaller than in the control (14.6+0.6 μm², p=0.009, and 14.9±0.8 μm², p=0.002, respectively). Analysis of variance confirmed the reliability of the dose-dependent effect of the RGDW peptide on the size of amorphous fibrin clusters (p<0.0001, Kruskal-Wallis test).

The observed decrease in the size of fibrin agglomerates under the influence of the RGDW peptide indicates indirectly a smaller size of platelet aggregates down to non-aggregated, single activated platelets, around which fibrin masses accumulate during the process of clot formation and remodeling.

### Mechanistic insights from the theoretical model

In agreement with the experiments, the theoretical plot in Fig. 7A predicts that (a) the final extent of clot contraction is independent of the RGDW concentration, (b) the rate of clot contraction slows as RGDW concentration increases, and (c) the lag time increases as RGDW concentration increases, which altogether validates the model.

(i) Fig. 7A shows that final extent of clot contraction is independent of RGDW concentration. Therefore, eventually all activated platelets are bound to the fibrin network and cause it to contract, although RGDW delays the process of platelets binding to fibrin.
(ii) Since fibrinogen-bound platelets form aggregates, the model result in Fig. 7A also implies that the force exerted by each platelet on the fibrin network is independent of whether it is individually bound to fibrin due to the presence of RGDW (Fig. 6B-D) or as part of an aggregate (Fig. 6A).
(iii) Fig. 7A shows that the rate of clot contraction (i.e., slope of the curve) decreases with increase in RGDW concentration. According to the model, the reason that RGDW delays clot contraction is that the rate at which platelets without fibrinogen (but with RGDW) bind to fibrin is much slower than the rate at which platelets with fibrinogen bind to fibrin. Figure S6 in the Supplemental Material, section 2, shows that if, on the contrary, the rate at which fibrinogen-bound platelets attached to fibrin was slower than platelets without fibrinogen then increasing RGDW would lead to faster contraction, which contradicts experimental observations.
(iv) Fig. 7B shows a model prediction of various platelet populations as a function of time for 0 and 200 μM RGDW. The magenta curve in Fig. 7B shows that a substantial population of platelets are bound to the RGDW peptide and this is why the increase in the population of platelets bound to fibrinogen (blue curves) and to fibrin (red curves) occurs slower and later.
(v) Figs. 7A and B show that the lag time is a consequence of positive feedback that results in the production of activated platelets due to stimulants released by other activated and aggregated platelets. The lag time for clot contraction increases as RGDW concentration increases in our model because (a) fewer fibrinogen-bound platelets are available to bind to fibrin and cause contraction, and (b) the rate of platelet activation is assumed to decrease as RGDW concentration increases.

**Figure 7.**
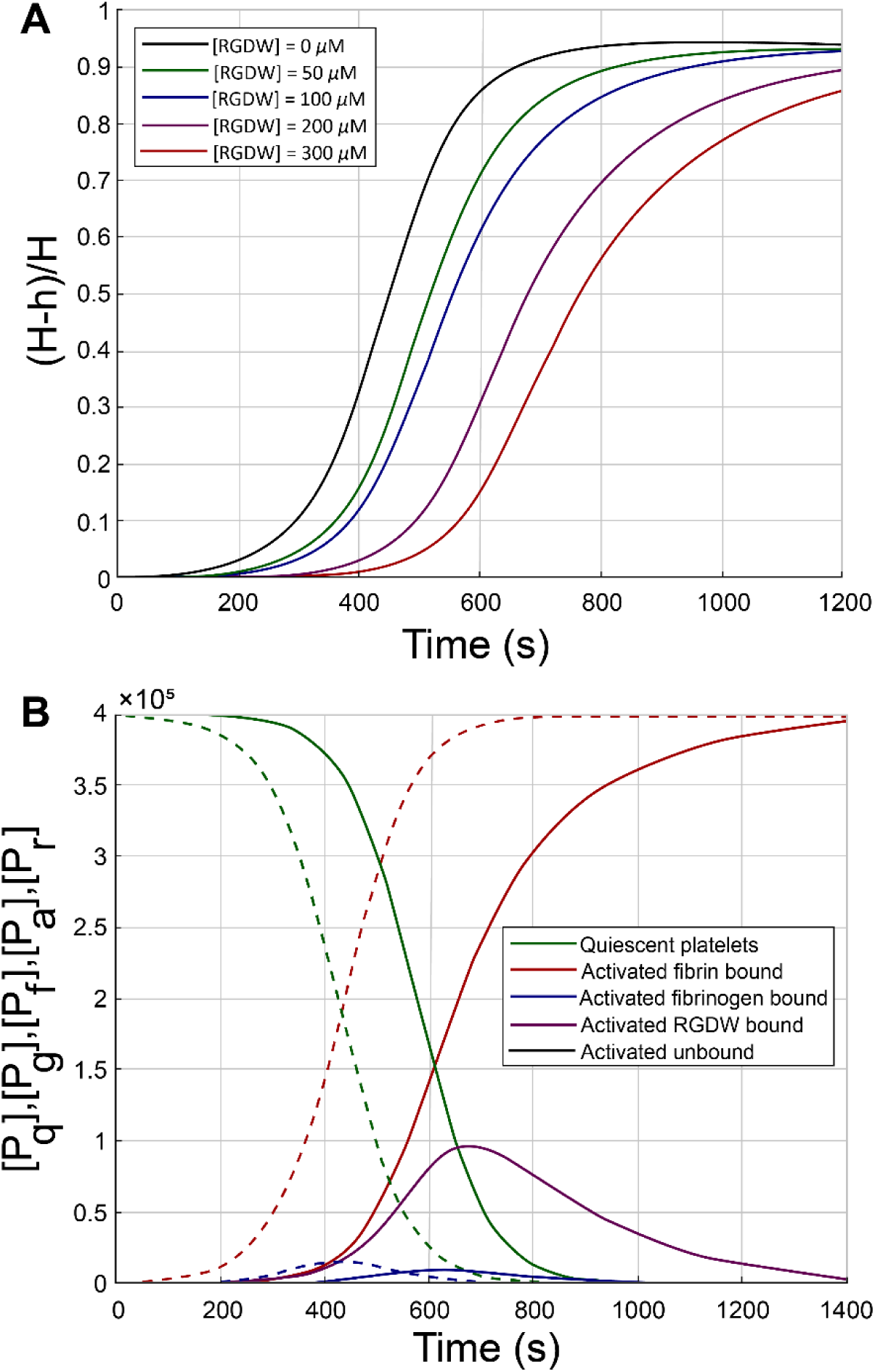
(**A**) Extent of clot contraction as a function of time as calculated from the theoretical model. *H* is the original area of the clot and *h(t)* is the current area. As the RGDW concentration increases contraction slows down and the lag time increases, in agreement with experiments. (**B**) Model prediction showing various platelet populations as a function of time in the absence (*dashed lines*) and presence (*solid lines*) of 200 μM RGDW. The magenta curve shows a large population of RGDW-bound platelets that eventually goes to zero at long times. The presence of RGDW prevents platelet aggregation and reduces the number of fibrin(ogen)-bound platelets attaching to fibrin, slowing contraction.

## Discussion

The study of platelet contractility has become increasingly important in recent years, due to emerging data on the critical role of clot contraction in both protective hemostasis and pathological thrombosis. In hemostasis, blood clot contraction is the final stage in the formation of a stable hemostatic plug.^2^ Clot contraction brings the wound edges closer together, thereby improving hemostasis and preventing wound infection.^4,5^ Compression of the fibrin network by activated platelets leads to its compaction and increased mechanical strength, which transforms the loose initial clot into a dense, impermeable plug resistant to blood flow pressure.^6^ During the formation and maturation of a mural thrombus, the contracted clot becomes less obstructive and less prone to fragmentation and rupture, i.e., contraction can reduce the risk of thromboembolism (e.g., pulmonary embolism and ischemic stroke).^7,8^ Conversely, impaired platelet contractility, leading to the formation of loose, unstable thrombi, may be a risk factor for thrombotic embolism. Contraction alters substantially the architecture of the blood clot, causing a spatial redistribution of components, with accumulation of the fibrin-platelet network at the periphery and densely packed red blood cells in the clot core.^33^ The fibrin network is compacted and becomes less porous, and red blood cells undergo compressive deformation with the formation of polyhedra called polyhedrocytes.^2,34^ Contracted blood clots have increased sensitivity to internal fibrinolysis and are more resistant to external (therapeutic) fibrinolysis.^9,10,35^ Contraction of clots and thrombi, which determines their density and rigidity, also affects the efficacy of mechanical thrombectomy.^36,37^

Platelet contractility is highly variable and can be impaired in both hereditary and acquired thrombocytopathy. Several hereditary diseases are known that directly affect key elements of the platelet contractile machinery. A classic example of such pathological conditions is macrothrombocytopenia syndrome with impaired clot contraction and hemorrhagic diathesis, caused by mutations in the *MYH9* gene, which encodes the heavy chain of non-muscle myosin IIA.^38,39^ Glanzmann’s thrombasthenia, caused by a qualitative defect and/or deficiency of platelet integrin αIIbβ3, is also accompanied by impaired platelet aggregation and blood clot contraction.^40,41^ In clinical practice, contractile function disorders are common because of acquired thrombocytopathy accompanying inflammatory conditions and (pro)thrombotic conditions that cause chronic platelet activation in the bloodstream and secondary platelet contractile dysfunction due to their energy depletion.^42,43^

It remains unclear how clinically used antiplatelet medications affect platelet contractility and blood clot contraction. The most commonly used antiplatelet drugs include inhibitors of the P2Y12 purine receptor (ticlopidine, clopidogrel, ticagrelor, prasugrel, cangrelor)^44^ or cyclooxygenase (acetylsalicylic acid).^45,46^ Integrin αIIbβ3 antagonists, such as abciximab, eptifibatide, and tirofiban, which block integrin binding to fibrinogen and inhibit platelet aggregation, are widely used.^18,27,47^ The effect of these drugs on blood clot contraction and thrombi remains unclear, as their effect on the interaction of integrin αIIbβ3 with fibrin and on mechanotransduction processes is unknown. Meanwhile, the most common and dangerous side effect of antiplatelet drugs is bleeding, and it is quite possible that one of the mechanisms of this complication is the suppression of platelet contractility, which normally promotes hemostasis.^38,48,49,50^ This assumption is supported by our data showing dose-dependent inhibitory effects of abciximab on the extent of blood clot contraction and corresponding reduction in clot stiffness (Figs. S2-S4).

Beyond its practical significance, a scientific problem is the functional relationship between two fundamental processes mediated by a single receptor (αIIbβ3), i.e., primary platelet aggregation and subsequent clot contraction. Although both processes are critical for hemostasis and thrombosis, it was unclear whether aggregation is essential for subsequent contraction and how exactly inhibiting platelet aggregation affects platelet contractility and contraction of blood clots and thrombi, which is the topic of this paper.

To study the relationship between platelet aggregation and contractility, we selectively inhibited platelet aggregation using the synthetic peptide, RGDW, which disrupts the interaction of platelet integrin αIIbβ3 with fibrinogen without affecting its binding to fibrin.^31^ The main result of our studies is that inhibition of platelet aggregation by the RGDW peptide (Fig. 1) slows the rate of contraction without affecting the final degree of clot compression in both whole blood (Fig. 2A) and plasma (Fig. 3A). In addition to contraction kinetics, inhibition of platelet aggregation alters the mechanical properties of the clot (Figs. 4D, 5D). Inhibition of platelet aggregation by the RGDW peptide alters the structure of the fibrin network, namely, it becomes more homogeneous (Fig. 6A-D) with a decrease in the size of amorphous fibrin agglomerates around platelets (Fig. 6E), reflecting the predominance of individual activated platelets over larger aggregates formed in the absence of RGDW. The seemingly paradoxical increase in whole blood clot stiffness could be due to the more uniform fibrin network in the presence of RGDW, arising from smaller, dispersed platelet/fibrin agglomerates. This homogeneous platelet-fibrin meshwork may interact differently with entrapped erythrocytes, perhaps compressing them more evenly, thereby leading to mechanically stiffer whole blood clots compared to the PRP clots.

To understand the mechanisms by which primary fibrinogen binding to activated platelets influences subsequent clot contraction, we used a theoretical model. The model comprises a system of ordinary differential equations that track various sub-populations of platelets as a function of time. It assumes irreversible binding of platelets to fibrin and fibrinogen, but reversible binding of the RGDW peptide to platelets, and shows (both analytically and through computations) how this leads to the final extent of clot contraction being independent of the RGDW concentration. The fact that final extent of clot contraction is independent of RGDW concentration also implies that fibrinogen-mediated platelet aggregation (which is inhibited by RGDW) has no direct effect on the contractile force exerted by platelets on fibrin. If the contractile force depended inversely on the surface area of platelet aggregates, this would lead to the result that the contractile force and hence clot contraction should decrease as the size of the platelet aggregates increases (see Supplemental Material section 2.4.1 and Figure S11), but this is not seen in the experiment.

The model also points to the mechanism behind the dose-dependent slowdown of clot contraction by RGDW as being the slower rate of attachment to fibrin of platelets without fibrinogen than those with fibrinogen. These platelet-bound fibrinogen molecules would quickly get converted to monomeric fibrin by thrombin and rapidly enter the fibrin network through polymerization, if there were no RGDW. At higher RGDW concentration, more platelets are prevented from binding to fibrinogen, which is incorporated into the fibrin network upon conversion to fibrin monomer. This is how RGDW slows the rate of clot contraction and increases lag time. It is suggested in the Supplemental Material that another possible reason for the increased lag time with increasing RGDW concentration is the disruption of positive feedback for platelet activation which is likely helped by fibrinogen-mediated aggregation of platelets. RGDW prevents aggregation of platelets, which might allow individual activated platelets to diffuse away, preventing their contact with quiescent platelets and reducing the overall rate of activation.

The practical significance of the results obtained lies in the potential for developing agents, perhaps peptide mimetics, with a more controlled and selective antithrombotic effect, possibly with an improved safety profile in terms of bleeding risk. Peptide mimetics with antiplatelet activity based on the RGD motif are being actively sought worldwide.^51–54^ However, in most cases, these peptides are tested primarily for their effect on platelet aggregation, and their effect on clot contraction is not considered. Our data show that antiplatelet agents, such as the RGDW peptide, impair platelet aggregation but do not inhibit clot contraction and instead modulate the kinetics of the process and alter clot structure and mechanical properties. Compounds with such properties that also prevent thrombosis without impairing hemostasis have been developed recently^55,56^ and may represent a new generation of future antithrombotic medications capable of not simply "switching platelets off" but selectively modulating their diverse functions.

In addition to their potential pharmacological implications, the results obtained have important theoretical significance, since they provide a new perception of the functional plasticity of platelets in hemostasis and thrombosis. It can be argued that aggregation and contraction are not simply two sequential events but connected processes. Our data demonstrate that the biological role of platelets can be realized within different time frames: from the rapid formation of a dense plug during massive injury to a slower, controlled clot compaction and remodeling, which may be important for maintaining the balance between hemostatic and thrombotic reactions in blood flow. In some biological and clinical settings, the kinetics of clot contraction or how fast it occurs can be more important than the final degree of shrinkage because the physiological consequences occur during the evolution of the clot, not just in its end state. First, timing determines whether a vessel remains open or becomes occluded and the blood flow stops. If contraction occurs rapidly, the clot quickly becomes dense and the hemostatic clot or thrombus may become less permeable and more resistant to external fibrinolysis or thrombolysis. These changes that occur within minutes to hours can influence whether blood flow is preserved or blocked and the surrounding tissue experiences a very different period of ischemia.

In conclusion, this study provides insight into the modulating role of fibrinogen-mediated platelet aggregation during platelet-driven clot contraction. Using the synthetic peptide RGDW, it was demonstrated that platelet aggregation and contractility can be pharmacologically decoupled. Modulation of the interaction of integrin αIIbβ3 with fibrinogen was found not to suppress clot contraction but to significantly alter its kinetics and the structural and mechanical properties of the fibrin network.

## Abbreviations

RGD: Arg-Gly-Asp;
RGDW: Arg-Gly-Asp-Trp;
AGDV: Ala-Gly-Asp-Val;
PRP: platelet-rich plasma;
ATP: adenosine triphosphate;
TEG: Thromboelastography;
ADP: adenosine diphosphate
TxA_2_: thromboxane A2;
SEM: scanning electron microscopy;
TRAP: thrombin receptor activating peptide

## Authorship Contributions

**Alina I. Khabirova:** Methodology, Investigation, Writing – original draft. **Rafael R. Khismatullin:** Methodology, Investigation. **Shakhnoza M. Saliakhutdinova:** Methodology, Investigation. **Natalia G. Evtugina:** Methodology, Investigation. **Lorena Buitrago:** Conceptualization, Methodology, Writing – review & editing**. Prashant K. Purohit:** Methodology, Investigation, Writing – original draft, Funding acquisition. **Rustem I. Litvinov:** Writing – original draft, Writing – review & editing, Supervision, Conceptualization. **John W. Weisel:** Conceptualization, Writing – review & editing, Supervision, Funding acquisition.

## Acknowledgements

We would like to thank Professor Barry Coller for fruitful discussions. The authors thank HemaCore Ltd. (Russia) for providing the Thrombodynamics Analyzer System used in this study.

## Supplemental Material

Section 1: Supplemental Methods; Figures S1-S4; Tables S1–S10

Section 2: Theoretical Model; Figures S5-S12; Table S11

