## Supplemental Methods for "The Role of Fibrinogen-Mediated Platelet Aggregation in Subsequent Platelet-Driven Blood Clot Contraction"

### Section 1. Supplemental methods, figures and tables

#### 1. Supplemental methods

*Fractionation of blood obtained from healthy donors*

Venous blood was obtained by venipuncture using VACUETTE® vacutainers (Greiner Bio-One, Austria) containing 3.2% sodium citrate in a 9:1 ratio by volume. Platelet-rich plasma (PRP) was obtained from citrated blood by centrifugation at 200g for 10 minutes at room temperature. Drawing blood from healthy donors over 18 years of age was approved by the Ethics Committee of Kazan (Volga Region) Federal University (Protocol No. 27 dated December 28, 2020). All whole blood and PRP samples were used within 2 hours. A total of 31 blood samples from independent donors were obtained and used in the study.

*Platelet aggregation in the presence of different concentrations of the RGDW peptide*

Platelet aggregation was studied in PRP using a Biola 220LA light aggregometer (Russia) in the absence or presence of the RGDW peptide (at concentrations of 50, 100, 200, and 300 μM) after pre-incubation for 5 minutes at 37°C. The peptide RGDW was synthesized by the Rockefeller University Proteomics Resource Center. Platelet aggregation was induced by the addition of thrombin receptor activating peptide 6 (TRAP 6, Sigma-Aldrich, USA) at a final concentration of 10 μM. The result was assessed as the final degree of platelet aggregation (percentage).

*Clot contraction in the presence of different concentrations of the RGDW peptide*

The kinetics of clot contraction in whole blood or PRP was studied by optically tracking clot size over time using a Thrombodynamics Analyzer (HemaCore, Russia). Before use, a transparent plastic cuvette (12 mm×7 mm×1 mm) was rinsed with either a 4% vol/vol Triton X-100 solution (for whole blood) or a 1% Pluronic F-127 solution (Millipore Sigma, USA) (for PRP) to prevent clot adhesion to the cuvette walls. In a separate plastic tube, clotting of blood or PRP (200 μl) was initiated by adding calcium chloride and human thrombin at final concentrations of 2 mM and 1 U/ml, respectively. 80 μl of the activated blood or PRP sample were rapidly transferred to a measuring cuvette preheated to 37°C in the recorder thermostat. 2 mM $CaCl_2$ was added to promote Factor XIIIa activation, fibrin cross-linking, and retention of red blood cells within the clot [Blood, 2016; 127: 149-59]. Photographic recording of the clot during compression was performed automatically every 15 seconds for 20 minutes. Digital clot images were used to build a contraction kinetic curve, which was used to calculate four parameters: 1) the final degree of contraction (the difference between the initial and final clot sizes as a percentage of the initial); 2) the lag period (the time from the addition of thrombin to the onset of clot shrinkage); 3) the slope of the kinetic curve (the tangent of the angle formed by the tangent line emanating from the end point of the lag period), characterizing the rate of clot contraction; and 4) the area under the kinetic curve, reflecting the integrated intensity of the process, which is determined collectively by the lag period, velocity, and final degree of clot contraction. Clot contraction assays were performed in the absence or presence of RGDW peptide (at concentrations of 50, 100, 200, and 300 μM) pre-incubated with whole blood or PRP for 5 min at 37°C.

*Thromboelastography of whole blood and PRP in the presence of different concentrations of the RGDW peptide*

Thromboelastography was performed using a Haemoscope TEG® 5000 device (Haemonetics, USA). Whole blood or PRP was incubated in the absence or presence of the RGDW peptide (at concentrations of 50, 100, 200, and 300 μM) with a kaolin suspension (Standard Technology, Russia) at a final concentration of 0.01 mg/ml for 5 minutes at 37°C. The kaolin-activated sample (340 μl) was transferred to the measuring cuvette of the Thromboelastograph, which was preliminarily supplemented with 20 μl of 360 mM $CaCl_2$ to a final concentration of 20 mM. The following parameters were automatically determined from the thromboelastogram (TEG): *R* (min) – reaction time (time from initiation of coagulation with calcium chloride to the onset of fibrin formation by the divergence of TEG lines to 2 mm), *K* (min) – time of increase in TEG amplitude from 2 mm to 20 mm (characteristic of the initial rate of fibrin polymerization); $\alpha^{o}$ – TEG tilt angle, characterizing the rate of clot formation (similar in value to *K*); *MA* (mm) is the maximum amplitude reflecting the maximum elasticity of the clot (*G*), which was calculated using the formula $G$ (Pa) = [500 × *MA*/(100 - *MA*)], recommended by the manufacturer of the TEG® 5000 instrument. It should be noted that the parameters *K* and the angle $\alpha$ both characterize the kinetics of clot formation, but their changes have opposite directions: a shortening of the *K* time is associated with an increase in the angle $\alpha$ and vice versa.

*Scanning electron microscopy of clots formed in PRP in the presence of different concentrations of the RGDW peptide*

Clots formed in PRP in the absence or presence of RGDW peptide (at concentrations of 50, 100, 200, and 300 μM) were allowed to contract for 20 min and then thoroughly washed with a physiological buffer to remove proteins, fixed with 2% glutaraldehyde, incubated for 90 min at room temperature, and stored at 4–6°C until use. Before scanning electron microscopy, the clots were washed with 50 mM phosphate buffer containing 150 mM NaCl (pH 7.4) and dehydrated with ethanol in ascending concentrations from 30% to 100% (vol/vol). Next, the samples were incubated with hexamethyldisilazane and air-dried at room temperature overnight. The dried samples were sputter-coated with gold-palladium (Polaron e5100 or Quorum Q150T ES devices) and then visualized on a scanning electron microscope (Merlin, Carl Zeiss, Germany).

*Statistical analysis*

Statistical analysis was performed using the GraphPad Prism 8.0.1 software package. Normality of data distribution was assessed using the D'Agostino–Pearson, Kolmogorov–Smirnov, and Shapiro–Wilk tests. Pairwise statistical differences were assessed using the Student's *t*-test (parametric) or Mann-Whitney U test (non-parametric). One-way ANOVA and Kruskal-Wallis test was used for multivariate analysis. Results are presented as the mean and standard error. The statistical significance level was 95% ($p<0.05$).

#### 2. Supplemental figures and tables

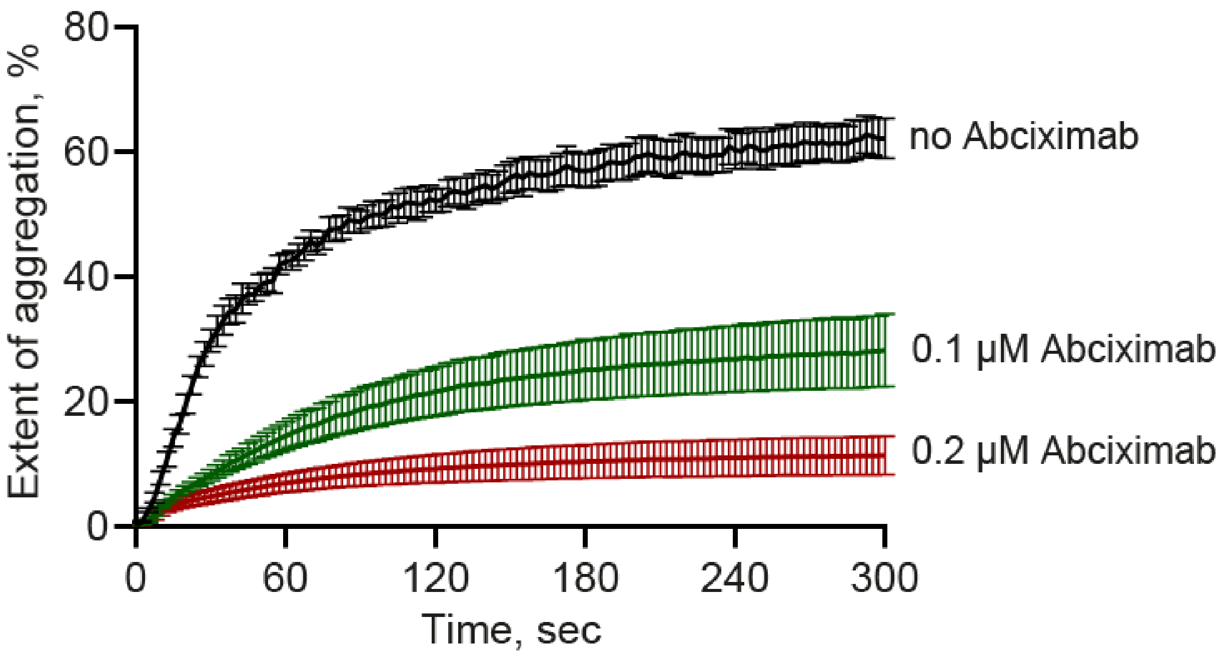


**Figure S1.** Platelet aggregation in PRP induced by 50 μM TRAP in the absence (*black curve*) and presence of the abciximab at various concentrations of 0.1 μM and 0.2 μM. The results are presented as the mean ± SEM (n=7). For numerical data see Table S2.

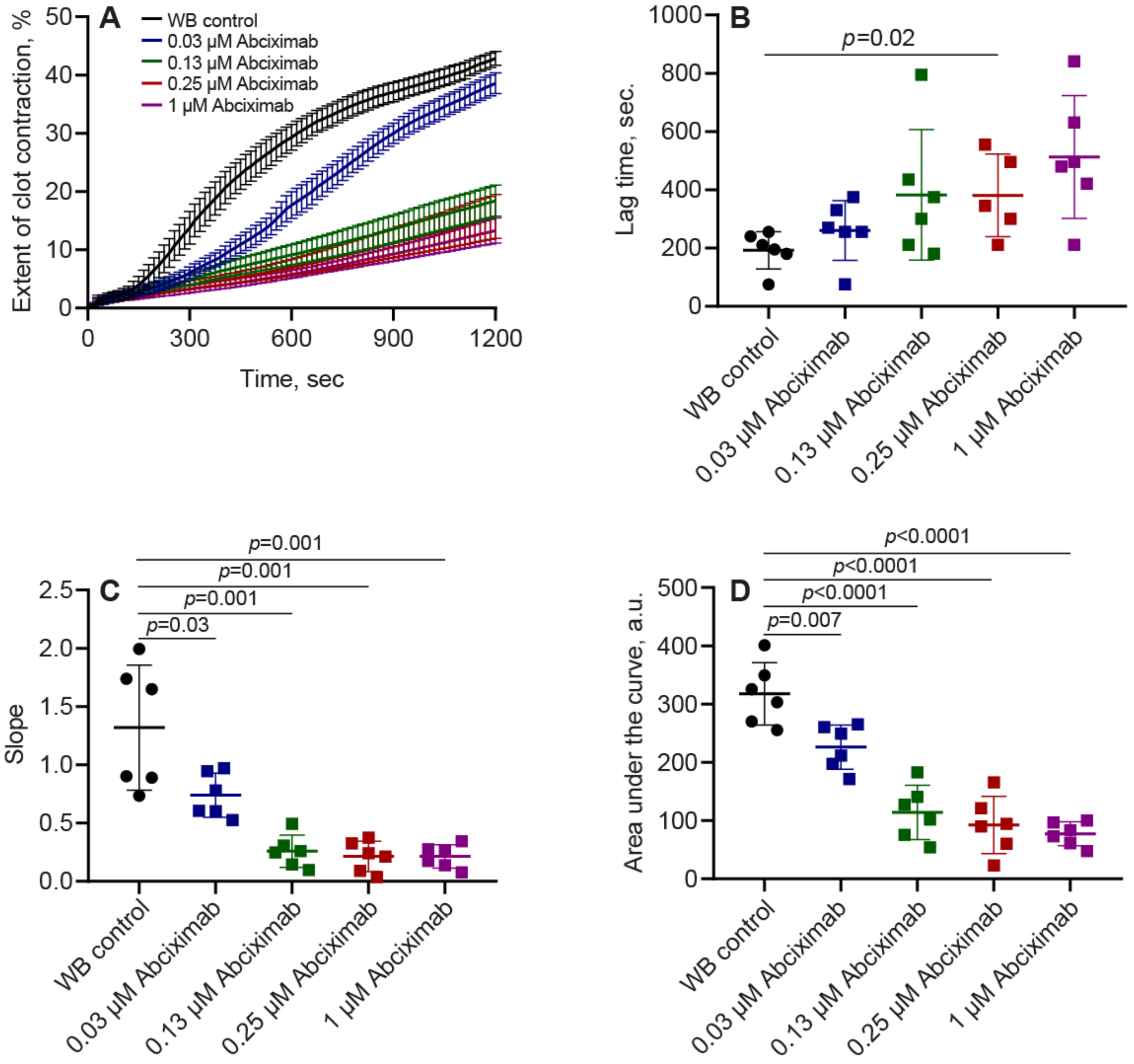


**Figure S2.** Contraction kinetic curves (**A**) and contraction parameters (**B-D**) of **clots formed in whole blood** (WB) in the absence and presence of abciximab at various concentrations of 0.03 μM, 0.13 μM, 0.25 μM, 1 μM. The results are presented as the mean ± SEM (n=6). The *p*-values reflect significant differences with the control, determined with the unpaired Student's *t*-test. For numerical date see Table S5.

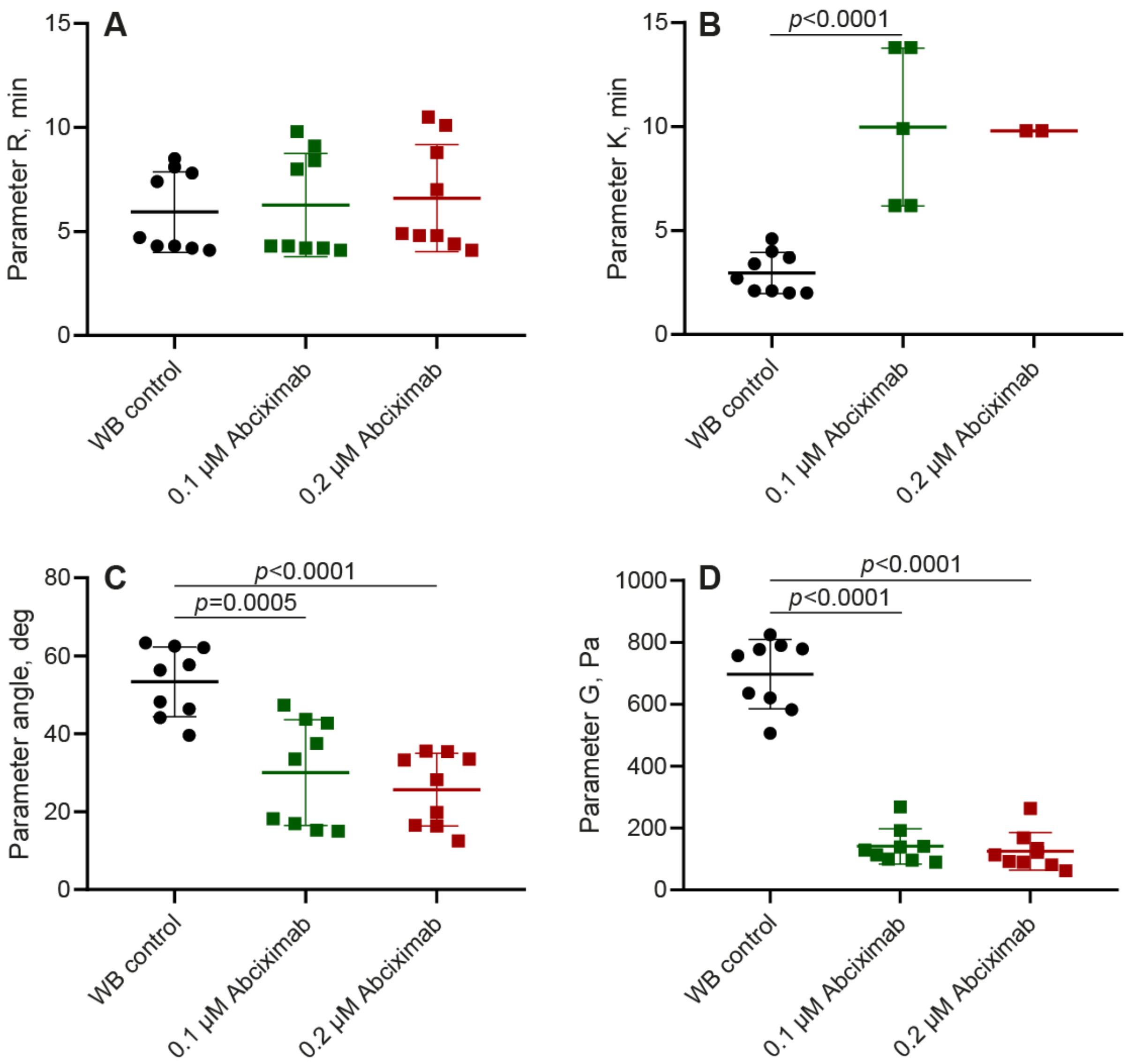


**Figure S3.** Parameters of TEG obtained in **whole blood** in the absence and presence of abciximab at the concentrations of 0.1 µM and 0.2 µM. In B, the parameter K could be determined only for the samples with MA>20 mm. The results are presented as the mean ± SEM (n=9). The *p*-values reflect significant differences with control determined with the Student's *t* test or Mann-Whitney U test. For numerical date see Table S8.

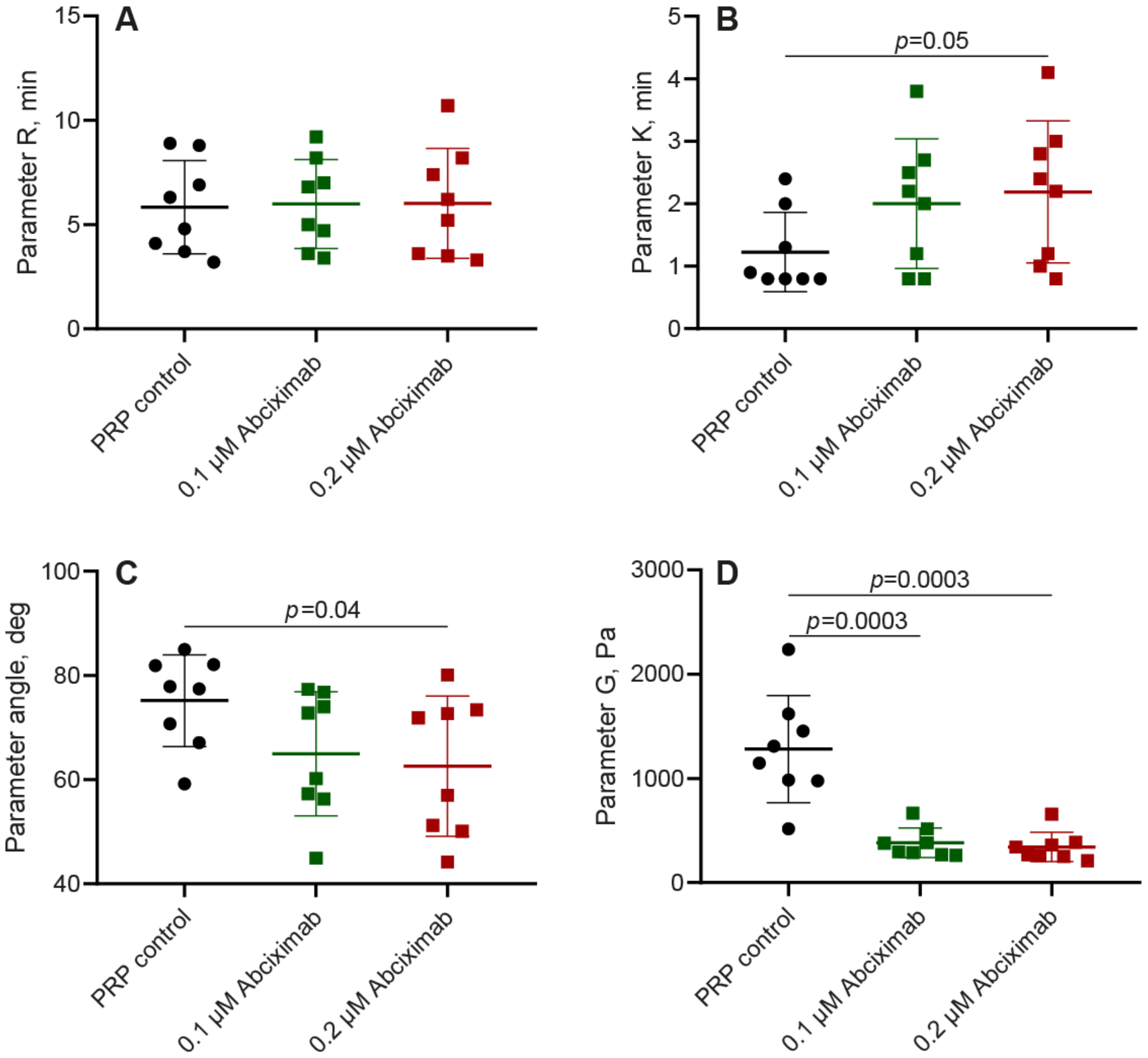


**Figure S4.** Parameters of TEG obtained in **PRP** in the absence and presence of abciximab at various concentrations of 0.1 µM and 0.2 µM. The results are presented as the mean ± SEM (n=8). The *p*-values reflect significant differences with control determined with the Student's *t* test or Mann-Whitney U test. For numerical date see Table S9.

Table S1. The final degree of platelet aggregation induced by 10 µM TRAP in the presence of the RGDW peptide at concentrations of 50 µM, 100 µM, 200 µM, 300 µM

| Final degree of aggregation (%) | *PRP+ TRAP (control)* | *PRP+50 µM RGDW + TRAP* | *PRP+100 µM RGDW + TRAP* | *PRP+200 µM RGDW + TRAP* | *PRP+300 µM RGDW + TRAP* |
|---|---|---|---|---|---|
| Mean ± SEM (n=3) | 58±4 | 51±6 | 39±4 | 21±4 | 10±2 |
| Paired *t*-test (comparison with control) | | 0.4 | ***0.02*** | ***0.02*** | ***0.02*** |
| *One-way ANOVA* | ***<0.0001*** | | | | |

Table S2. The final degree of platelet aggregation induced by 10 µM TRAP in the presence of abciximab at concentrations of 0.1 µM and 0.2 µM.

| Final degree of aggregation (%) | *PRP+TRAP (control)* | *PRP+0.1 µM abciximab + TRAP* | *PRP+ 0.1 µM abciximab + TRAP* |
|---|---|---|---|
| Mean ± SEM (n=7) | 62±3 | 28±6 | 11±3 |
| Paired *t*-test (comparison with control) | | ***0.002*** | ***<0.0001*** |
| *One-way ANOVA* | ***<0.0001*** | | |

Table S3. Parameters of **whole blood** clot contraction in the absence and presence of the RGDW peptide at concentrations of 50 µM, 100 µM, 200 µM, 300 µM.

| | *Extent of clot contraction, %* | *Lag time, sec.* | *Slope* | *Area under the curve, a.u.* |
|---|---|---|---|---|
| Whole blood control (n=8) | 46±2 | 180±17 | 1.68±0.18 | 350±20 |
| WB+50 µM RGDW (n=7) | 46±1 | 221±16 | 1.62±0.14 | 335±13 |
| WB+100 µM RGDW (n=8) | 46±1 | 268±21 | 1.51±0.14 | 308±8 |
| WB+200 µM RGDW (n=8) | 46±1 | 313±15 | 1.58±0.14 | 293±8 |
| WB +300 µM RGDW (n=8) | 43±1 | 377±17 | 1.19±0.08 | 243±9 |
| *One-way ANOVA* | 0.3 | ***<0.0001*** | 0.1 | ***<0.0001*** |

Table S4. Parameters of **PRP** clot contraction in the absence and presence of the RGDW peptide at concentrations of 50 µM, 100 µM, 200 µM, 300 µM.

| | *Extent of clot contraction, %* | *Lag time, sec.* | *Slope* | *Area under the curve, a.u.* |
|---|---|---|---|---|
| PRP control (n=9) | 90±2 | 122±9 | 2.72±0.10 | 822±38 |
| PRP+50 µM RGDW (n=9) | 91±1 | 190±27 | 2.52±0.08 | 780±36 |
| PRP+100 µM RGDW (n=6) | 89±3 | 213±74 | 2.15±0.10 | 708±38 |
| PRP+200 µM RGDW (n=9) | 90±2 | 315±51 | 1.86±0.08 | 612±75 |
| PRP+300 µM RGDW (n=6) | 89±3 | 398±78 | 1.42±0.03 | 498±77 |
| *One-way ANOVA* | p=0.2 | ***p<0.0001*** | ***<0.0001*** | ***p<0.0001*** |

Table S5. Parameters of **whole blood** clot contraction in the absence and presence of the abciximab at concentrations of 0.03 µM, 0.13 µM, 0.25 µM, 1 µM.

| | *Extent of clot contraction, %* | *Lag time, sec.* | *Slope* | *Area under the curve, a.u.* |
|---|---|---|---|---|
| Whole blood control | 44±1 | 193±26 | 1.32±0.22 | 318±22 |
| WB+0.03 µM abciximab | 40±2 | 260±41 | 0.74±0.08 | 226±16 |
| WB+0.13 µM abciximab | 18±3 | 383±91 | 0.26±0.06 | 114±19 |
| WB +0.25 µM abciximab | 15±4 | 381±63 | 0.21±0.05 | 92±20 |
| WB +1 µM abciximab | 12±3 | 513±86 | 0.21±0.04 | 77±8 |
| *One-way ANOVA (n=6)* | ***<0.0001*** | ***0.01*** | ***<0.0001*** | ***<0.0001*** |

Table S6. Parameters of thromboelastography (TEG) of **whole blood** in the presence and absence of peptide at concentrations of 50 µM, 100 µM, 200 µM, 300 µM

| *Parameters of TEG* | *R, min* | *K, min* | *Angle, degree* | *G, pascal* |
|---|---|---|---|---|
| Whole blood control | 4.5±0.5 | 2.1±0.3 | 62±3 | 622±57 |
| WB+50 µM RGDW | 4.6±0.5 | 2.3±0.2 | 60±2 | 612±58 |
| WB+100 µM RGDW | 4.3±0.6 | 2.4±0.5 | 60±5 | 613±87 |
| WB+200 µM RGDW | 4.5±0.4 | 2.2±0.2 | 62±2 | 800±51 |
| WB +300 µM RGDW | 4.2±0.4 | 3.4±0.2 | 53±1 | 794±80 |
| *One-way ANOVA (n=5)* | 0.5 | 0.06 | 0.1 | ***0.03*** |

Table S7. Parameters of thromboelastography (TEG) of **PRP** in the presence and absence of peptide at concentrations of 50 μM, 100 μM, 200 μM, 300 μM

| *Parameters of TEG* | *R, min* | *K, min* | *Angle, degree* | *G, pascal* |
|---|---|---|---|---|
| PRP control | 3.7±0.3 | 0.8±0 | 81±1 | 1433±125 |
| PRP +50 μM RGDW | 3.2±0.3 | 0.8±0 | 82±1 | 1327±119 |
| PRP +100 μM RGDW | 3.5±0.4 | 0.8±0 | 81±1 | 1341±203 |
| PRP +200 μM RGDW | 3.5±0.2 | 0.8±0 | 79±0.3 | 1459±173 |
| PRP +300 μM RGDW | 3.5±0.3 | 1.1±0.1 | 76±1 | 1626±173 |
| *One-way ANOVA (n=5)* | 0.3 | ***0.05*** | ***<0.0001*** | 0.3 |

Table S8. Parameters of thromboelastography (TEG) of **whole blood** in the presence and absence of abciximab at concentrations of 0.1 μM and 0.2 μM

| *Parameters of TEG* | *R, min* | *K*, min* | *Angle, degree* | *G (pascal)* |
|---|---|---|---|---|
| Whole blood control | 5.9±0.6 | 2.9±0.3 | 53.4±3.0 | 697±37 |
| WB +0.1 μM Abciximab | 6.2±0.8 | 10.0±1.7 (n=5) | 30.0±4.5 | 141±19 |
| WB +0.2 μM Abciximab | 6.6±0.9 | 9.8±0 (n=2) | 25.7±3.1 | 125±20 |
| *One-way ANOVA/ Kruskal-Wallis test (n=9)* | 0.6 | ***<0.0001*** | ***<0.0001*** | ***<0.0001*** |

*The parameter K could be determined only for the samples with MA>20 mm.

Table S9. Parameters of thromboelastography (TEG) of **PRP** in the presence and absence of abciximab at concentrations of 0.1 µM and 0.2 µM

| *Parameters of TEG* | *R, min* | *K, min* | *Angle, degree* | *G (pascal)* |
|---|---|---|---|---|
| PRP control | 5.8±0.8 | 1.2±0.2 | 75.2±3.1 | 1281±181 |
| PRP+0.1 µM Abciximab | 5.9±0.8 | 2.0±0.4 | 65.0±4.2 | 383±50 |
| PRP+0.2 µM Abciximab | 6.0±0.9 | 2.2±0.4 | 62.6±4.8 | 343±50 |
| One-way ANOVA/ Kruskal-Wallis test (n=8) | 0.9 | 0.1 | 0.9 | ***0.0007*** |

Table S10. Fibrin agglomerate area (µm$^2$) in PRP clots in the presence and absence of peptide at concentrations of 50 µM, 100 µM, 200 µM, and 300 µM

| | *Area of fibrin agglomerates, µm$^2$* | *Unpaired t-test (comparison with control)* |
|---|---|---|
| PRP control | 20.7±1.2 | |
| PRP +50 µM RGDW | 14.2±1.4 | 0.0003 |
| PRP +100 µM RGDW | 12.6±0.8 | <0.0001 |
| PRP +200 µM RGDW | 14.6±0.6 | 0.009 |
| PRP +300 µM RGDW | 14.9±0.8 | 0.002 |
| *One-way ANOVA (Kruskal-Wallis test) (n=5)* | <0.0001 | |

### Section 2. Theoretical Model

#### 1 A very brief mechanistic picture of platelet activation

Initially, platelets are quiescent or resting. Platelets can get activated due to (a) biochemical signals, (b) shear stress, and (c) adhesion to collagen or von Willebrand factor. The main physiological stimulants that activate platelets are thrombim, ADP, and thromboxane A2 (TxA2). Activated platelets show (a) cytoskeletal changes, (b) secretion of high and low molecular weight compounds, (c) high adhesion to fibrinogen and fibrin, and (d) contractility. The integrins on the membrane of activated platelets bind readily to many ligands and fibrinogen which has two binding sites for the integrin. As a result, activated platelets can aggregate with fibrinogen molecule acting as a bridge between two platelets. Activated and aggregated platelets release ADP and TxA2 which activate other quiescent platelets leading to a positive feedback. The integrin binding sites in fibrin and fibrinogen are different, and so is their specificty and binding strength. Fibrinogen bound to platelets gets converted to fibrin by thrombin and could get incorporated into fibrin fibers rapidly.

#### 2 Kinetic model of platelet activation

We want to take account of the above facts in our model of platelet activation and contractility. Let $P_0$ be the total number of platelets in a vessel of one microliter. $P_q(t)$ is the number of quiescent platelets at time $t$. We set $t = 0$ when contact of thrombin with platelets occurs. Fibrin formation also starts at this time, but platelet activation happens faster than fibrin polymerization. The time scale of platelet activation and clot formation is a few minutes. This observation will inform our choice of rate constants. The number of activated platelets is $P_0 - P_q(t)$. Activated platelets are divided into three populations. $P_f(t)$ is the number of activated platelets bound to fibrin, $P_g(t)$ is the number of activated platelets bound to fibrinogen, and $P_a(t)$ is the number of activated platelets bound neither to fibrin nor fibrinogen. We know that $P_0 = P_q + P_f + P_g + P_a$. If platelets gets activated and aggregated they cause other platelets to get activated due to positive feedback. Hence, the number of activated platelets increases as

$$\frac{d}{dt}(P_f + P_g + P_a) = k_0(P_f + P_g + P_a)(P_0 - P_f - P_g - P_a), \tag{1}$$

where $k_0$ is a constant. We re-write the equation above as:

$$\frac{dP_q}{dt} = -k_0(P_0 - P_q)P_q. \tag{2}$$

We will assume that platelets once bound to fibrin do not detach. Activated platelets that are bound to fibrinogen and those that are not bound to fibrinogen can both bind to fibrin. We will assume different rate constants for these processes. We will assume that each activated platelet occupies $a$ binding sites on fibrin. $F$ is the number of fibrin binding sites available to activated platelets in a one microliter vessel. Then, the rate of increase of $P_f(t)$ is

$$\frac{dP_f}{dt} = k_1 P_a(F - aP_f) + k_2 P_g(F - aP_f), \tag{3}$$

where $k_1$ and $k_2$ are different rate constants. We suspect from experiments that $k_2 > k_1$, i.e., the rate $k_2$ at which fibrinogen bound platelets bind to fibrin is higher than the rate $k_1$ at which activated platelets without fibrinogen bind to fibrin. The fibrinogen bound to platelets gets converted to fibrin monomer by thrombin and then it becomes part of the fibrin polymerization process which proceeds rapidly in parallel with platelet activation. Also, aggregation of platelets due to platelet-to-platelet contact and via bound fibrinogen speeds up secondary activation of quiescent platelets due to secretion of activating stimuli. For these reasons activated platelets with bound fibrinogen likely have greater affinity to bind to fibrin than those platelets which do not have bound fibrinogen, leading to $k_2 > k_1$. Finally, the rate of change of $P_g(t)$ is:

$$\frac{dP_g}{dt} = -k_2 P_g(F - aP_f) + k_3 P_a. \tag{4}$$

The first term on the RHS says that fibrinogen bound platelets get consumed when they bind to fibrin fibers. The constant $k_3$ in the second term on the RHS accounts for the rate at which activated platelets with no fibrin or fibrinogen converts to platelets bound to fibrinogen, assuming that excess fibrinogen is present in the vessel. We know from experiments that eventually all platelets get activated and bound to fibrin. Do our governing equations predict this outcome? To answer this question we will look for steady states of the above three ODEs and check if they allow a solution in which $P_f \to P_0$ as $t \to \infty$. We need to solve three equations for the three unknown steady state values $P_f$, $P_a$, $P_g$, given some $P_0$, $F$, $a$, etc.

$$\begin{aligned} 0 &= -k_0(P_0 - P_q)P_q, & (5)\\ 0 &= k_1 P_a(F - aP_f) + k_2 P_g(F - aP_f), & (6)\\ 0 &= -k_2 P_g(F - aP_f) + k_3 P_a. & (7) \end{aligned}$$

From the first equation $P_q = P_0$ is a steady state. In this steady state $P_a = P_f = P_g = 0$ for all time. This steady state occurs only if no platelets are activated at $t = 0$. The first equation gives another steady state in which $P_q = 0$, i.e., no quiescent platelets. Assuming $F - aP_f > 0$, because the number of fibrin binding sites is in excess of what all platelets could cover, the second and third equations imply:

$$\begin{aligned} k_1 P_a + k_2 P_g &= 0, & (8)\\ k_3 P_a - k_2 P_g(F - aP_f) &= 0. & (9) \end{aligned}$$

Solving eqn. (8) for $P_a$ and using $P_f = P_0 - P_a - P_g$ we can rewrite eqn. (9) as:

$$k_2 P_g \left[ \frac{k_3}{k_1} + F - a[P_0 + P_g(\frac{k_2}{k_1} - 1)] \right] = 0. \tag{10}$$

$P_g = 0$ is indeed a solution of the above. Then, from eqn. (8) $P_a = 0$, and since $P_f = P_0 - P_g - P_a$ we must have $P_f = P_0$. This is the desired steady state. Another solution is $P_g = \frac{1}{\frac{k_2}{k_1} - 1}\left[\frac{k_3/k_1 + F}{a} - P_0\right]$. Assuming that there are fibrin binding sites in excess of those that can be covered by $P_0$ platelets, the above can be positive only if $k_2 > k_1$, that is, the rate governing the binding of fibrinogen bound platelets attaching to fibrin is larger than the rate governing activated platelets without fibrinogen attaching to fibrin. Therefore, we will assume $k_2 > k_1$ throughout. Later, we will give a stronger reason for assuming $k_2 > k_1$ which is rooted in how the model prediction compares with experimental observation.

We need to determine some parameters. A crucial parameter is $a$ which is the number of fibrin binding sites occupied by a single activated platelet. We will estimate $a$ as follows. Each platelet is $3\mu$m in diameter and it contains about 80,000 integrins, 40,000 on each side of the disc. A fibrin fiber is about 100nm in diameter. Let us assume that all integrins in a rectangle 100nm $\times 3\mu$m in size on the platelet surface are attached to a fibrin fiber which is 100nm in diameter. The number of integrins in this rectangle is roughly $\frac{40000 \times 0.1\mu m \times 3\mu m}{(\pi/4) \times 3\mu m \times 3\mu m} \approx 1698$. Therefore, $a$ is likely to be a few thousand. Another way to estimate $a$ is as follows. A single platelet integrin transmits forces in the range of 12 to 54pN during adhesion and spreading. While initial adhesion can occur at lower forces, platelet contraction and clot stabilization can involve higher tensions, with some single molecule studies giving bond rupture forces as high as $90 - 100$pN [1] . A single activated platelet can apply an average maximum contractile force of approximately 29 to 34 nN (nanonewtons) on a fibrin network, with, in some cases, forces exceeding 100 nN. These platelets can form adhesions stronger than 70 nN and can bend or stretch fibrin fibers by hundreds of nanometers during clot contraction [2]. Therefore, we estimate $a$ to be between 200 to 3000. We take $a = 5000$. We take $P_0 = 400,000$ per microliter, as in platelet rich plasma. We take $F = 2 \times 10^{10}$, so $F >> aP_0$, i.e., the number of binding sites for activated platelets is much larger than 400,000 platelets can occupy. We need to assume values for the constants $k_0, k_1, k_2, k_3$. We do this by trial and error as we solve the three ODEs eqns. (2), (3) and (4) to see how various platelet populations change in time. The right values of the constants $k_0$, $k_1$, $k_2$, $k_3$ are those that give good agreement with experimental results on clot contraction. We used the following constants (all in SI units not given below) for figure S2.

$$
\begin{aligned}
P_0 &= 400,000, \quad F = 2 \times 10^{10}, \quad a = 5000, \quad k_0 = 5 \times 10^{-8}, \quad k_1 = 5 \times 10^{-14}, \\
k_2 &= 5 \times 10^{-12}, \quad k_3 = 1.3 \times 10^{12}. \qquad (11)
\end{aligned}
$$

Notice that in figure S2 the number of fibrinogen bound platelets remains low for all time while that of fibrin bound platelets monotonically increases. This is a consequence of $k_2 > k_1$. The number of activated unbound platelets is very small because they bind either to fibrin or fibrinogen quickly after activation. This a consequnce of the large $k_3$ value chosen. The rate constants are chosen in such a way as fibrinogen bound platelets rapidly attach to fibrin. The population of activated platelets with fibrinogen increases in the beginning, but as time progresses it eventually goes to zero since the platelets end up attached to fibrin, as required by the steady state.

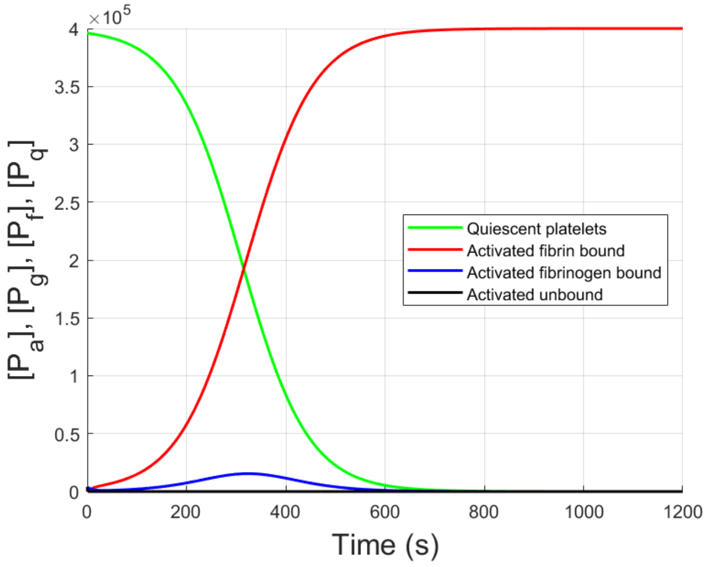


Figure S5: Numbers of quiescent and activated platelets as a function of time. We assume $P_0 = 400,000$ per microliter and $F = 2 \times 10^{10}$. Rate constants are chosen such that the number of fibrin bound platelets plateaus around 600s.

### 3 Effect of RGDW peptide

RGDW is a small peptide that competes with fibrinogen to bind to the integrins in activated platelets. When RGDW has blocked the integrins then the platelet cannot bind to fibrinogen, but it can still bind to fibrin. Therefore, RGDW bound platelets cannot aggregate and they bind individually to fibrin. A schematic showing the effects of RGDW appears in figure S3. RGDW can unbind from the integrins and then the activated platelet can bind to fibrinogen and fibrin. To take account of the effects of RGDW we will now have RGDW bound platelets as another population, $P_r$, in addition to $P_q$, $P_a$, $P_g$ and $P_f$. As before, $P_0 = P_q + P_a + P_g + P_f + P_r$. Let the number of RGDW peptides in our one microliter vessel be $R$ and let $b$ be the number of peptides required to block a platelet from binding to fibrinogen. Then, the rate at which $P_r$ evolves is:

$$\frac{dP_r}{dt} = k_4(R - bP_r)P_a - k_5P_r - k_1P_r(F - aP_f). \tag{12}$$

The first term accounts for the encounters of free floating RGDW and activated platelets, the second term accounts for RGDW unbinding from the platelets and the third term accounts for the RGDW bound platelets attaching to fibrin. The evolution equations for the other

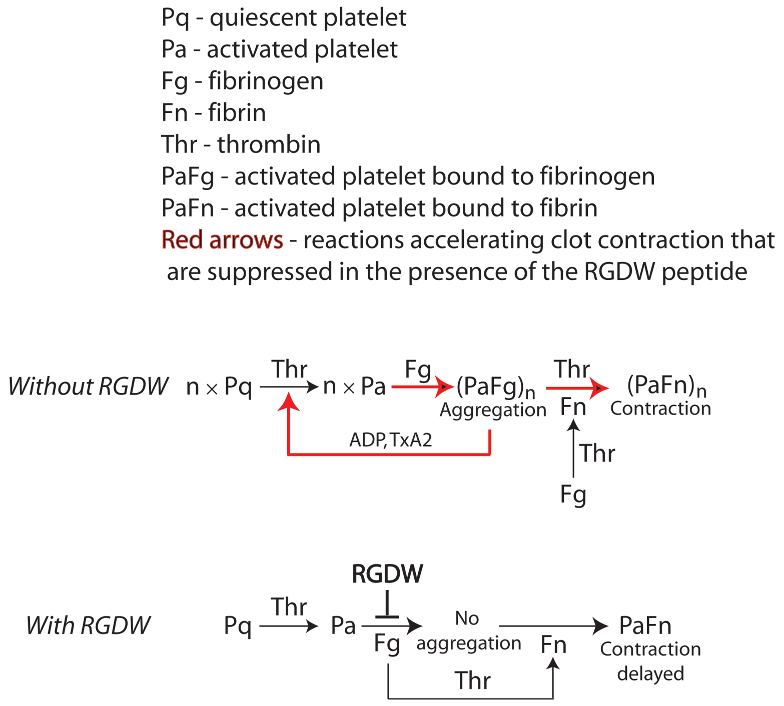


Figure S6: A schematic diagram showing how RGDW suppresses certain reactions in clot contraction caused by platelets.

four populations of platelets are now modified to:

$$\frac{dP_q}{dt} = -k_0 P_q(P_0 - P_q), \tag{13}$$

$$\frac{dP_f}{dt} = k_1(P_a + P_r)(F - aP_f) + k_2 P_g(F - aP_f), \tag{14}$$

$$\frac{dP_g}{dt} = -k_2 P_g(F - aP_f) + k_3 P_a, \tag{15}$$

$$P_a = P_0 - P_f - P_g - P_r - P_q. \tag{16}$$

In eqn. (14) we have assumed that activated platelets with or without RGDW bind to fibrin with the same rate constant $k_1$. We have assumed, as before, that once a platelet is bound to fibrinogen or fibrin it remains attached to them. The fibrinogen bound platelets can be incorporated into fibrin fibers after cleaving by thrombin. Only those activated platelets that have neither fibrinogen nor fibrin bound to them can bind RGDW. Again, we look for steady states to show that as $t \to \infty$, $P_f \to P_0$, i.e., all platelets in the vessel are activated and bound to fibrin, eventually. At steady state the equations to be solved for $P_q$, $P_a$, $P_g$, $P_f$ and $P_r$ are:

$$0 = -k_0 P_q(P_0 - P_q), \tag{17}$$

$$0 = k_1(P_a + P_r)(F - aP_f) + k_2 P_g(F - aP_f), \tag{18}$$

$$0 = -k_2 P_g(F - aP_f) + k_3 P_a, \tag{19}$$

$$0 = k_4(R - bP_r)P_a - k_5 P_r - k_1 P_r(F - aP_f), \tag{20}$$

$$P_a = P_0 - P_f - P_g - P_r - P_q. \tag{21}$$

As before, the first equation gives $P_q = 0$ (we neglect the trivial case $P_q = P_0$). From the second and third equations we find that both $P_a$ and $P_r$ are proportional to $P_g$. When, we substitute for $P_a$ and $P_r$ in terms of $P_g$ into the fourth equation we get $P_g = 0$ as a solution. Hence, it follows that $P_a = 0$ and $P_r = 0$. Then, the fifth equation gives $P_f = P_0$ in steady state, irrespective of the amount of RGDW peptide present. Therefore, irrespective of the amount of RGDW, all platelets in our vessel eventually bind to fibrin and exert contractile force on it. If each platelet exerts the same force on fibrin irrespective of whether it is an individual or part of an aggregate then the final extent of clot contraction will also be the same, independent of the amount of RGDW present.

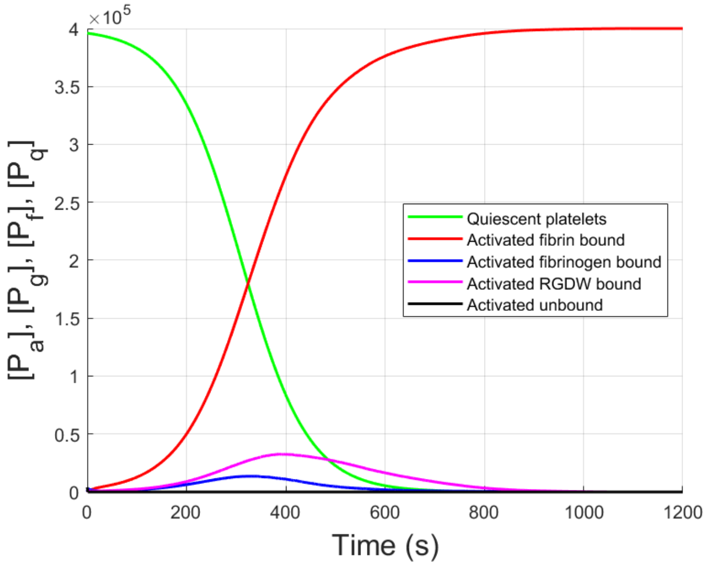


Figure S7: Numbers of quiescent and activated platelets as a function of time. We assume $P_0 = 400,000$ per microliter and $F = 2 \times 10^{10}$. RGDW concentration is taken to be $50\mu$M. In contrast to figure S 2, t he p opulation o f a ctivated u nbound p latelets $P_a(t)$ r emains very small while that of RGDW bound platelets $P_r$ is higher than fibrinogen b ound p latelets $P_g$ at all times. Since $k_2 > k_1$ and $k_1$ multiplies $P_r$ while $k_2$ multiples $P_g$, the speed of clot contraction is reduced in the presence of RGDW.

Again, we need to estimate some parameters. A crucial parameter is $b$, the number of RGDW peptides needed to block binding of a platelet to fibrinogen. S ince a n activated platelet contains about 80,000 integrins on its surface and each integrin has one RGDW binding site we take $b = 10^5$. We used this and the following parameters and integrated eqns. (12), (13), (14), (15), (16). The initial conditions were:

$$P_q(0) = 0.99P_0, \quad P_a(0) = 0.01P_0, \quad P_g(0) = P_f(0) = P_r(0) = 0. \tag{22}$$

For figure S4 we plot various platelet populations as a function of time while holding fixed $R = 3 \times 10^{13}$ ($50\mu$M works out to $3 \times 10^{13}$ RGDW peptides in one microliter). The key observation in figure S4 is that the number of RGDW bound platelets is larger than the number of fibrinogen bound platelets at all times; this slows down clot contraction since the rate at which platelets without fibrinogen attach to fibrin is slower than the rate at which platelets with fibrinogen attach to fibrin, i.e., $k_1 < k_2$. In experiments $R$ varies from 50$\mu$M to 300$\mu$M. So, we kept initial conditions fixed and varied $R$ and plotted the population of fibrin bound platelets as a function of time because these are responsible for clot contraction. The result is shown in figure S5. As RGDW concentration increases the rate of increase of fibrin bound platelets slows down and the lag time increases slightly. This will result in slower contraction and longer lag times, as seen in experiment. The parameters used to

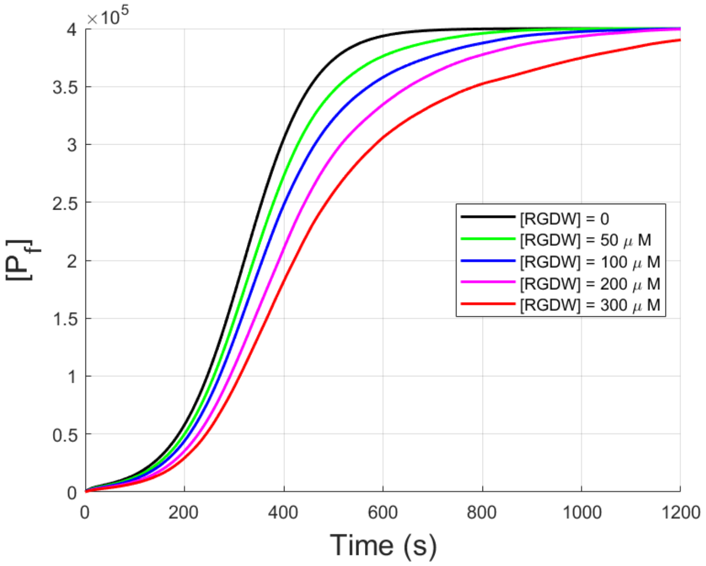


Figure S8: Number of fibrin bound platelets as a function of time and RGDW concentration. The lag increases as RGDW concentration increases, but it is not exactly the same as that seen in experiment. However, the trend observed in the kinetic curves is very similar to that in experiment.

make figures S4 and S5 are:

$$\begin{aligned} P_0 &= 400,000, \quad F = 2 \times 10^{10}, \quad a = 5000, \quad k_0 = 3.7 \times 10^{-8}, \quad k_1 = 5 \times 10^{-14}, \\ k_2 &= 5 \times 10^{-12}, \quad k_3 = 1.3 \times 10^{12}, \quad b = 10^5, \quad k_4 = 1 \times 10^{-2}, \quad k_5 = 5 \times 10^{-3}. \end{aligned} \tag{23}$$

To confirm that $k_1 < k_2$ is necessary to replicate experimental trends we plot the number of fibrin bound platelets as a function of time with $k_1 = 5 \times 10^{14}$ and $k_2 = 5 \times 10^{12} < k_1$ in figure S6. This time the trend is exactly opposite of that seen in experiment, viz., as RGDW concentration increases rate of increase of fibrin bound platelets speeds up. Therefore, we rule out the possibility that $k_1 > k_2$.

### 4 Clot contraction

To study clot contraction, we begin with a rectangular clot that is thin. In experiments the clot is 12mm×7mm×1mm. Platelets are all over this clot and when they contract the rectangle shrinks, as in the cuvette in the thromboimager used. Let the undeformed area of

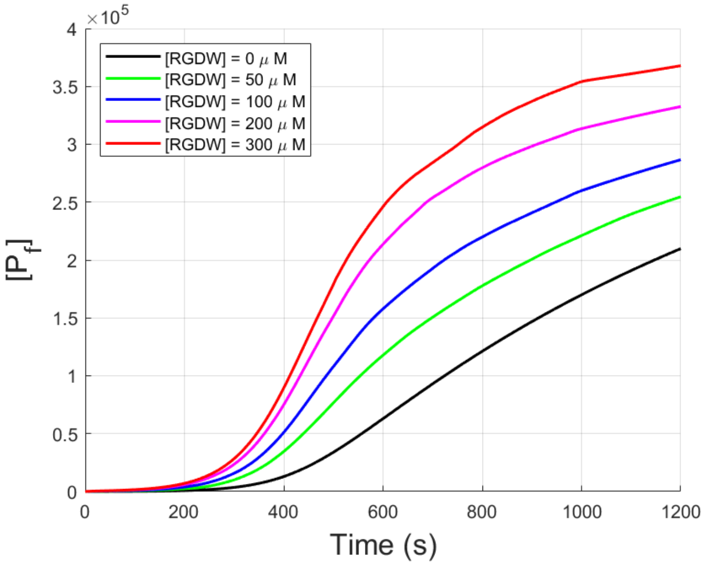


Figure S9: Number of fibrin bound platelets as a function of time and RGDW concentration when $k_1 > k_2$. The trend is exactly opposite of that seen in experiments. Therefore, we rule out the possibility that $k_1 > k_2$.

the clot be $H$ and the deformed area be $h$, then assuming a linear relation between stress in the clot and area strain we write

$$\sigma(t) = B(1 - \frac{h(t)}{H}) - \sigma_p(t), \tag{24}$$

where $B$ is a modulus of the clot material and $\sigma_p(t)$ is the hydrostatic tensile stress due to activated platelets attached to fibrin. We have taken compressive stresses to be positive and tensile stresses negative in writing the expression above. We will assume $\sigma_p(t)$ is proportional to $P_f(t)$, i.e., as soon as an activated platelet binds to fibrin it exerts its maximum contractile force on the fibrin immediately. Of course, this may not be true and a platelet will take some time to reach its maximum contractile force, but we will assume instantaneous activation for simplicity. As the clot contracts liquid leaves through the edges of the clot so that the liquid flux is proportional to $\frac{dh}{dt}$. By Darcy's law this flux must be proportional to the pressure gradient at the edges. We assume that all the pressure drop at the edges occurs over small distance $\delta << \sqrt{h}$. Then, by Darcy's law

$$\frac{dh}{dt} = \frac{k}{\eta}\frac{\sigma(t)}{\delta} = \frac{Bk}{\eta\delta}(1 - \frac{h}{H}) - \frac{k\sigma_p(t)}{\eta\delta}, \tag{25}$$

where $k$ is a Darcy constant and $\eta$ the viscosity of liquid. The above can be re-written as:

$$\frac{dh}{dt} + \frac{Bk}{H\eta\delta}h = \frac{Bk}{\eta\delta} - \frac{k\sigma_p(t)}{\eta\delta}. \tag{26}$$

In eqn. (25) we take $\sigma_p(t) = dP_f(t)$ where $d$ is a constant. We will take $B$ to be on the order of KPa. Each platelet produces about 35nN of force and the area of cross-section is

1$\mu$L/12mm, which gives $d = 0.42$ Pa. We will take $\delta$ to be about 100$\mu$m. The ODE eqn. (25) is combined with the other ODEs that determine the evolution of platelet populations to compute clot contraction. The initial conditions are the same as in eqn. (22) together with $h(0) = H$. We will plot $\frac{H-h(t)}{H}$ where $h(t)$ is the current area of the clot. The results of

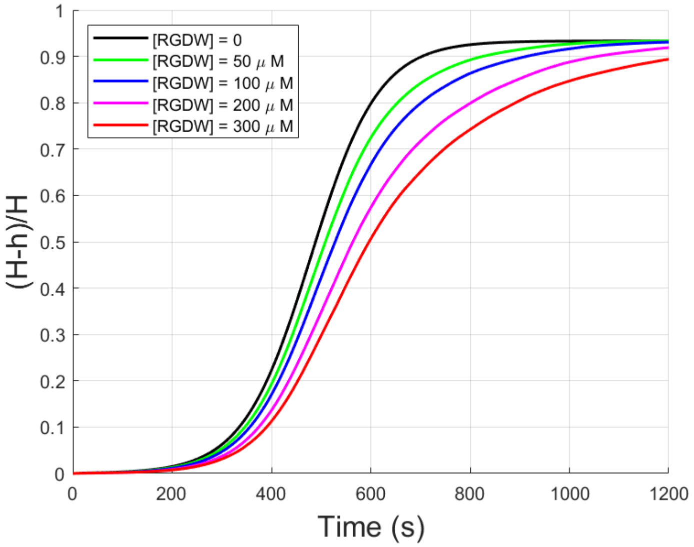


Figure S10: Extent of clot contraction as a function of RGDW concentration. The trends seen in experiment are reproduced. We do not see differences in lag time with RGDW concentration as clear as in experiment, but the shape of the kinetic curves is captured.

integrating eqns. (12), (13), (14, (15), (16) and (25) are shown in figure S7. The parameters are the same as in eqn. (29), except $P_q(0) = 0.999P_0$ for this calculation. The new parameters that enter this calculation are given below:

$$k = 2.6 \times 10^{-15} m^2, B = 180KPa, \delta = 100\mu m, \eta = 10^{-3} Pa.s, H = 1\mu L/12mm. \tag{27}$$

The modulus $B$ is a bit high because the clot densifies as it contracts and platelets contribute to stiffness. Figure S7 captures most trends seen in experiment, except the variation in lag time as a function of RGDW concentration. The lag time does increase in figure S7 as RGDW concentration increases, but not by the same amount as in experiment. We address the lag time in the next section.

#### 4.1 Does aggregation affect platelet contractile force?

A possibility exists that the force exerted by platelets depends on their surface area in contact with fibrin fibers since the binding sites for fibrin are on the surface, i.e., on platelet membranes. When platelets form an aggregate some of the surface area of each platelet is in contact with other platelets, so not all of it is available for contact with fibrin. We will account for this in a simple way by assuming that the surface area of a cluster of $N_c$ platelets scales as $N_c^{2/3}$. Therefore, the force per platelet in a cluster of $N_c$ platelets scales as

$N_c^{-1/3}$. If there are a total of $P_f$ platelets bound to fibrin then the stress caused by them is $\sigma_p = dN_c^{-1/3} P_f$. When $N_c = 1$ then each cluster is an individual platelet and we recover the model laid out previously. Now, we will take $P_f = 400,000$ fixed and plot the time trajectory of clot contraction for various values of $N_c$. We will integrate the ODE

$$\frac{dh}{dt} + \frac{Bk}{H\eta\delta} h = \frac{Bk}{\eta\delta} - \frac{kdN_c^{-1/3} P_f}{\eta\delta}, \tag{28}$$

where $P_f = 400,000$ is held fixed and all other parameters are as in eqn. (27). The result is plotted in figure S8. This figure suggests that platelet cluster size can have a big effect on the

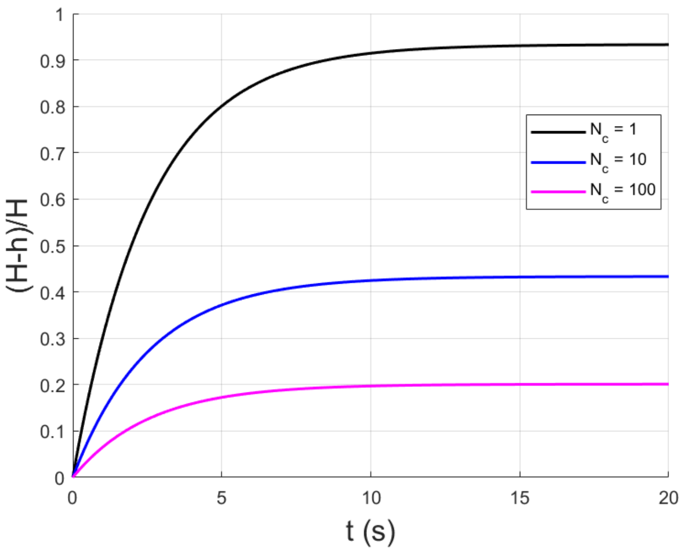


Figure S11: Extent of clot contraction as a function of time for different platelet cluster sizes. $P_f = 400,000$ is held fixed. As the cluster size is increased the extent of contraction decreases. This is not observed in experiments. Therefore, we reject the possibility that force per platelet depends on the size of the platelet aggregate.

extent of clot contraction. As the cluster size increases the total stress caused by platelets decreases, so the final extent of contraction also decreases. The slope of the contraction curve also decreases as cluster size increases. But, experiments show that the final extent of contraction is independent of RGDW concentration. RGDW affects aggregation and therefore platelet cluster size $N_c$. Therefore, it seems that RGDW does not affect the forces exerted by platelets on fibrin. Individual platelets and platelets in an aggregate exert the same force on fibrin fibers.

### 5 Lag time of clot contraction

The lag time of clot contraction seen in experiments is a manifestation of the sigmoid growth in the number of activated (including fibrin bound) platelets. The slope of the curve for $P_q(t)$ is nearly zero near $t = 0$ because the RHS of ODE eqn. (13) has $(P_0 - P_q)P_q$ which is small near $t = 0$. In other words, sigmoid growth is the direct result of the positive feedback mechanisms that cause an increase in the number of activated platelets. Therefore, the lag time must be governed by the parameter controlling positive feedback near $t = 0$ which is the product $k_0 P_q(0)(P_0 - P_q(0))$. It could be that RGDW modifies the constant $k_0$ in eqn. (13), but we instead consider the more likely possibility that RGDW modifies the number of (secondarily) activated platelets $P_0 - P_q(0)$ at $t = 0$. Our hypothesis is that the change in lag time due to the presence of RGDW can be captured if we assume that (a) the initial condition $P_q(0)$ is affected by RGDW ($P_0$ is fixed, so changing $P_0 - P_q(0)$ implies a change in $P_q(0)$), and (b) that observable contraction only starts when $P_f(t)$ reaches a minimum value. We will take the minimum $P_f = 4000$ for our one microliter vessel.

$$\begin{aligned} P_0 &= 400,000, \quad F = 2 \times 10^{10}, \quad a = 5000, \quad k_0 = 3.7 \times 10^{-8}, \quad k_1 = 5 \times 10^{-14}, \\ k_2 &= 5 \times 10^{-12}, \quad k_3 = 1.3 \times 10^{12}, \quad b = 10^5, \quad k_4 = 1 \times 10^{-2}, \quad k_5 = 5 \times 10^{-3}. \end{aligned} \tag{29}$$

With these parameters we used different initial conditions for $P_q(0)$ for each RGDW concentration to get lag times in the range of those observed in experiment. The lag times above

Table S11: Initial conditions on $P_q$ to reproduce lag times in experiment.

| **[RGDW]** | $P_q(0)$ | **Lag time** |
|---|---|---|
| 0$\mu$M | $0.99820P_0$ | 125.62s |
| 50$\mu$M | $0.99915P_0$ | 186.40s |
| 100$\mu$M | $0.99933P_0$ | 211.79s |
| 200$\mu$M | $0.99982P_0$ | 321.53s |
| 300$\mu$M | $0.99992P_0$ | 396.57s |

are those needed to reach $P_f = 4000$ starting from $P_f(0) = 0$. As the RGDW concentration increases, the product $P_q(0)(P_0 - P_q(0))$ decreases, and so the secondary rate of formation of activated platelets also decreases. This leads to longer lag times with increasing RGDW concentration. It could be that RGDW disrupts the contacts between activated platelets and nearby quiescent platelets reducing their ability to form aggregates slowing down secondary activation. A possible reason for this is that RGDW is a competitive inhibitor of platelet-fibrinogen interactions which promote aggregation and ultimately activation of platelets. Another possibility is that individaul RGDW bound platelets that do not aggregate can diffuse away so that they and the stimulants they produce do not come into contact with quiescent platelets quickly. A plot of clot contraction with these initial conditions appears in figure S9. This time we mostly capture the effects of increasing RGDW seen in experiments – (a) slow down in the rate of contraction, and (b) increase in lag time. We have already proved mathematically that the final extent of contraction is not affected by RGDW concentration.

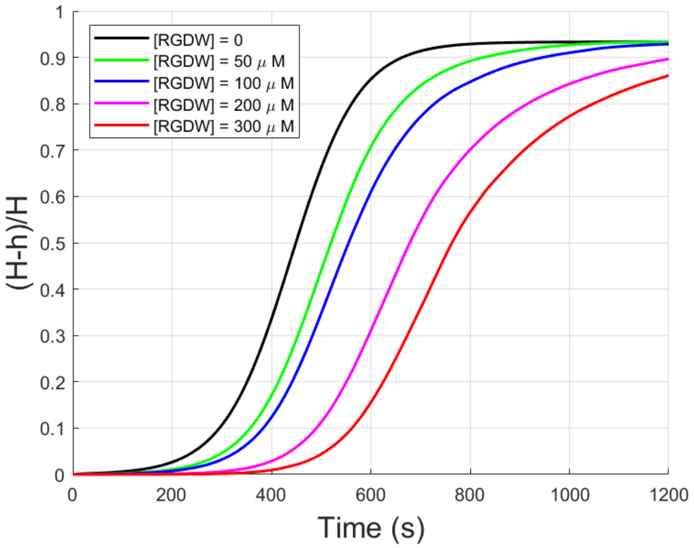


Figure S12: Extent of clot contraction as a function of RGDW concentration. We assumed that the initial condition is affected by RGDW concentration to get the correct lag times.

[2] Sun Y, Oshinowo O, Myers DR, Lam WA, Alexeev A. Resolving the missing link between single platelet force and clot contractile force. IScience. 2022 Jan 21;25(1).
